# Selective IL-27 production by intestinal regulatory T cells permits gut-specific regulation of Th17 immunity

**DOI:** 10.1101/2023.02.20.529261

**Authors:** Chia-Hao Lin, Cheng-Jang Wu, Sunglim Cho, Rasika Patkar, Ling-Li Lin, Mei-Chi Chen, Elisabeth Israelsson, Joanne Betts, Magdalena Niedzielska, Shefali A. Patel, Han G. Duong, Romana R. Gerner, Chia-Yun Hsu, Matthew Catley, Rose A. Maciewicz, Hiutung Chu, Manuela Raffatellu, John T. Chang, Li-Fan Lu

**Affiliations:** School of Biological Sciences, University of California, San Diego, La Jolla, California, CA, USA; Bioscience, Translational Science and Experimental Medicine, Research and Early Development, Respiratory & Immunology (R&I), BioPharmaceuticals R&D, AstraZeneca, Gothenburg, Sweden; Bioscience, Research and Early Development, Respiratory & Immunology (R&I), BioPharmaceuticals R&D, AstraZeneca, Gothenburg, Sweden; Department of Medicine, University of California, San Diego, La Jolla, CA, USA; Department of Medicine, Veterans Affairs San Diego Healthcare System, San Diego, CA, USA; Division of Host-Microbe Systems and Therapeutics, Department of Pediatrics, University of California, San Diego, La Jolla, CA, USA; Department of Pathology, University of California San Diego, La Jolla, CA, USA; Chiba University-UC San Diego Center for Mucosal Immunology, Allergy, and Vaccines (CU-UCSD cMAV), La Jolla, CA 92093, USA; Center for Microbiome Innovation, University of California, San Diego, La Jolla, CA, USA; Moores Cancer Center, University of California, San Diego, La Jolla, CA, USA

## Abstract

Regulatory T (Treg) cells are instrumental in establishing immunological tolerance. However, the precise effector mechanisms by which Treg cells control a specific type of immune response in a given tissue remains unresolved. By simultaneously studying Treg cells from different tissue origins under systemic autoimmunity, here we show that IL-27 is specifically produced by intestinal Treg cells to regulate Th17 immunity. Selectively increased intestinal Th17 responses in mice with Treg cell-specific IL-27 ablation led to exacerbated intestinal inflammation and colitis-associated cancer, but also helped protect against enteric bacterial infection. Furthermore, single-cell transcriptomic analysis has identified a CD83^+^TCF1^+^ Treg cell subset that is distinct from previously characterized intestinal Treg cell populations as the main IL-27 producers. Collectively, our study uncovers a novel Treg cell suppression mechanism crucial for controlling a specific type of immune response in a particular tissue, and provides further mechanistic insights into tissue-specific Treg cell-mediated immune regulation.

## Introduction

Regulatory T (Treg) cells control diverse types of immune responses and maintain immunological tolerance and tissue homeostasis ^1^. While the expression of Foxp3 as a master molecular regulator in Treg cells distinguishes them from other T cell lineages ^2–5^, it is now also well recognized that similar to other T cells that they regulate, Treg cells also come in different phenotypic and functional “flavors” ^6^. The acquisition of T helper (Th) cell lineage-specific transcription factors in Treg cells endows them with the capacities to control the corresponding immune responses in different inflammatory settings ^7–10^. Nevertheless, since transcription factors control the expression of a large number of genes, the precise effector mechanisms by which different Th-specific Treg cell subset control their respective type of T cell immunity have yet to be determined.

In addition to the Treg cell subsets that control different types of T cell immune responses, the presence of distinct Treg cell populations in many nonlymphoid tissues have now also been well appreciated ^11, 12^. Beyond exerting their immunoregulatory function to control local inflammation in a given anatomical site, these so-called tissue Treg cells were also shown to exhibit specific functional features to maintain corresponding organismal homeostasis. For example, during infection-inflicted lung damage or muscle and ischaemic stroke-induced brain injuries, Treg cells in lung, muscle, and brain are able to secret amphiregulin (Areg), a ligand of the epidermal growth factor receptor (EGFR) to facilitate tissue repair ^13–15^. In the skin, expression of Jagged 1 by Treg cells is crucial for efficient hair regeneration through driving Notch-dependent stem cell proliferation and differentiation ^16^. Moreover, in the adipose tissue, adenosine generated by CD73^+^ Treg cells was recently reported to promote adaptive thermogenesis by activating beige fat biogenesis ^17^. These studies have provided experimental evidence demonstrating the effector mechanisms underlying the non-immunological role of Treg cells in maintaining tissue homeostasis. On the other hand, like the aforementioned Th-specific Treg cells, how each tissue Treg cell population controls its corresponding local immune responses remains poorly characterized.

To date, only a handful of suppressor molecules have been implicated in tissue Treg cell-mediated immune regulation ^12^, the most well characterized of which is IL-10. Treg cell-specific IL-10 deletion did not result in the same degree of severe systemic autoimmunity that was shown with Foxp3 disruption or Treg-specific deletion of other known Treg cell-suppressor molecules like CTLA4 ^18–20^. Yet mice with IL-10 deleted specifically in Treg cells exhibited inflammation at multiple mucosal sites such as the skin, lung and colon, indicating that IL-10, albeit not a universal Treg suppressor molecule, was still commonly utilized by many tissue Treg cell subsets ^18^. Consistent with this notion, it was also suggested that Treg cells that reside in the visceral adipose tissue (VAT) mediate their anti-inflammatory effect by producing a large amount of IL-10 that would in turn act on neighboring adipocytes expressing the corresponding receptor ^21^. In addition, Treg cell-derived IL-10 also does not seem to regulate a specific type of immune response since mice harboring Treg cells incapable of producing IL-10 exhibited exacerbated Th1, Th2, and Th17-driven tissue pathology ^18, 22^. By employing an experimental system that permits simultaneous examination of multiple tissue Treg cell subsets during active suppression of systemic autoimmunity, here we identified distinct transcriptomic signatures in two different tissue Treg cell populations that could account for their respective control of local inflammation. In particular, we found that IL-27, a pluripotent cytokine recognized for its regulatory properties ^23^, is specifically induced in gut Treg cells under inflammatory conditions. Moreover, IL-27 derived from Treg cells but not from other known intestinal IL-27-producing cell populations is selectively needed for controlling Th17 responses in the gut-associated tissue. Loss of IL-27 expression by Treg cells led to exacerbated Th17-driven intestinal inflammation and colitis-associated cancer. On the other hand, enhanced Th17 responses in mice with Treg cell-specific IL-27 ablation could also help protect against enteric bacterial infection. Finally, unlike any other intestinal Treg cells that have been reported previously, single-cell transcriptomic analysis of intestinal Treg cells revealed a distinct CD83^+^TCF1^+^ Treg cell subset that does not express IL-10 but is responsible for IL-27 production particularly during intestinal inflammation. Together, our study uncovers a previously uncharacterized Treg cell suppression mechanism that is pivotal for controlling a specific type of immune response in a particular tissue and provides further mechanistic insights into tissue-specific Treg cell-mediated immune regulation.

## Results

### Identification of unique transcriptome signatures in tissue Treg cells during active suppression of local inflammatory responses

To date, most tissue Treg cell transcriptomic analyses were done by studying a specific population of tissue Treg cells isolated from mice under a defined disease condition (e.g. muscle Treg cells from a muscle injury model) ^11^, thus precluding stringent comparative bioinformatics analysis of Treg cell subsets in different tissues from the same mouse. In cases where different tissue Treg cells were isolated from the same animal, only unmanipulated and unchallenged mice were used. As it has become evident that, like conventional T (Tconv) cells, Treg cells differentiate into effector Treg cells upon activation and that both T cell receptor (TCR) activation and cytokine stimulation play critical roles in inducing Treg cell suppressor activity ^24, 25^, one could argue that much information about tissue-specific suppressor molecules involved in the active Treg cell-mediated suppressor program is still missing. To capture the dynamic gene expression profiles in tissue Treg cells involved in restricting ongoing autoimmune-mediated inflammation *in vivo*, we adapted and further modified a previously described experimental system allowing us to simultaneously assess multiple tissue Treg cells when they are actively controlling ongoing autoimmunity ^26^. In brief, multi-organ autoimmune inflammation was induced by the ablation of the entire Treg cell population upon diphtheria toxin (DT) administration in mice containing the coding sequence of a DT receptor (DTR) inserted into the 3’ untranslated region (UTR) of Foxp3 (*Foxp3^DTR^*) ^27^. Instead of transiently depleting Treg cells through the brief administration of DT as shown in the previous study ^26^, endogenous Treg cells were kept ablated by continuous injection of DT (**Fig. S1A**). The fatal consequence of DT-mediated Treg cell deletion was then rescued via the transfer of congenically marked Treg cells (**Fig. S1B, C** and data not shown). This way, we were able to obtain synchronized *in vivo* activated Treg cells for our transcriptomic study without the contamination of recent thymic Treg cell emigrants that might not be properly activated. Next, rather than waiting until the complete resolution of the autoimmune inflammation, Treg cells and Tconv cells from a secondary lymphoid tissue (i.e. spleen) as well as from the lung and small intestinal lamina propria (SI LP) were isolated from DT-treated *Foxp3^DTR^* mice or from control PBS-treated mice 10 days after Treg cell transfer for RNA-sequencing (RNA-seq) and comparative bioinformatics analysis. As shown in **Fig. S1D**, despite the fact that the DT-treated *Foxp3^DTR^* mice did eventually recover due to the presence of transferred Treg cells, at this time point, Tconv cells remained highly activated and expressed genes characteristic of the Th1, Th2 and Th17 subsets of effector T (Teff) cells.

Before investigating the impact of autoimmune inflammation on different tissue Treg cells, we first validated the results of tissue Treg cells at steady state. In agreement with previous studies ^12^, our analysis found genes that are preferentially upregulated in Treg cells from the non-lymphoid tissues, including transcription factors (e.g. *Irf4, Nfil3, Id2, Rorc* and *Fosl2*), cytokine receptors (e.g. *Il1rl1*), effector molecules (e.g. *Il10, Klrg1, Areg, Gzmb, Fgl2, Metrnl* and *Lrb4r1*) and co-inhibitory molecules (e.g. *Pdcd1* and *Lag3*). Similarly, we also detected genes that are known to be expressed at the lower level in tissue Treg cells such as *Id3, Tcf7, Bach2* and *Nrp1* (**Fig. 1A**). Nevertheless, despite a clear difference in gene expression observed in different tissue Treg cell subsets isolated from either DT-treated *Foxp3^DTR^* mice or PBS-treated control mice, the impact of ongoing autoimmune inflammation on the transcriptional profiles in Treg cells (and Tconv cells) was apparent (**Fig. 1B**). Moreover, consistent with the differences shown in the heatmap analysis, principal-component analysis (PCA) of the RNA-seq results revealed a high degree of resemblance in Treg cells (as well as corresponding Tconv cells) isolated from mice with DT treatment compared to the PBS-treated controls regardless of which tissues they came from (**Fig. 1C**, top). On the other hand, both Treg cells and Tconv cells in the same tissue also shared a considerable level of similarity in gene expression regardless of the presence or absence of an inflammatory condition (**Fig. 1C**, bottom). While certain genes such as Foxp3 are consistently highly expressed in Treg cells (data not shown), our RNA-seq study clearly demonstrated that both autoimmune inflammation and environmental factors greatly influence Treg cell gene expression profiles.

**Figure 1.**
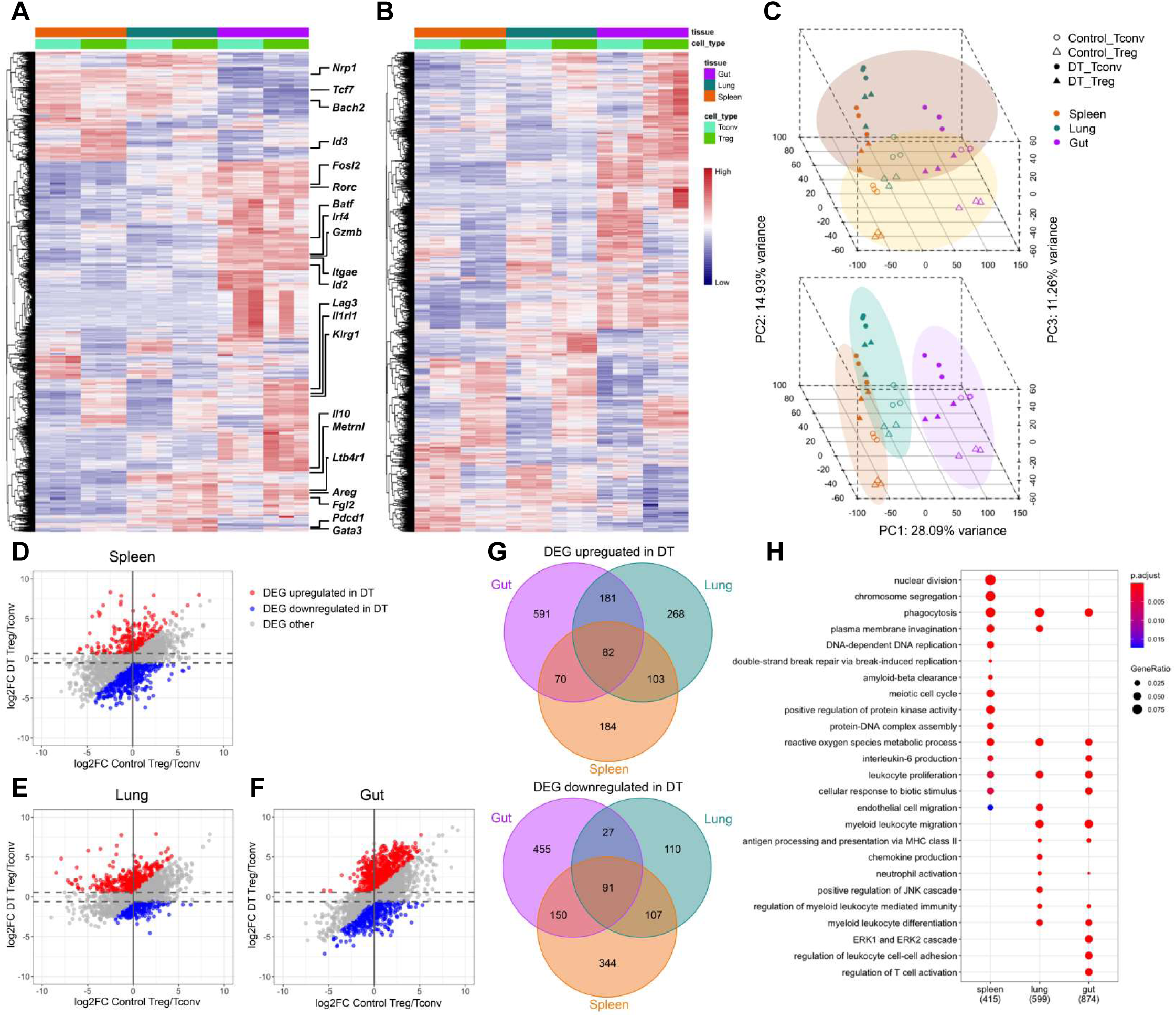
Transcriptomic analysis of tissue Treg cells during active suppression of local inflammatory responses. Heatmaps of top 10% of most variable genes in Treg cells isolated from indicated tissues in **(A)** control PBS-treated or **(B)** DT-treated *Foxp3^DTR^* mice 10 days after Treg cell transfer. **(C)** PCA of gene expression by different Treg and Tconv cell subsets. Different cell samples were grouped by treatment (top) or anatomical location (bottom). Scatter plots depicting log2 fold changes of gene expression in **(D)** spleen, **(E)** lung and **(F)** gut (SI LP) Treg cells over Tconv cells in DT-treated mice versus those in PBS-treated controls. Genes that are upregulated under DT-treated condition (Red; false discovery rate (FDR) < 5% and the value of log2 fold change (log2FC) more than 0.585 (1.5 fold) and at least 0.585 higher than the value of log2FC under PBS-treated control condition); Genes that are downregulated under DT-treated condition (Blue; FDR < 5% and the value of log2FC less than −0.585 and at least 0.585 lower than the value of log2FC under PBS-treated control condition). **(G)** Venn diagrams demonstrating genes up- or down-regulated in different tissue-specific Treg cells from DT-treated mice. Numbers represent gene numbers. **(H)** Dot plot of Gene Ontology (GO) term enrichment analysis of differentially expressed genes (DEG) upregulated in tissue Treg cell subsets. Colors indicate the p values from Fisher’s exact test, and dots size is proportional to the number of DEG in a given biological process.

Next, to gain further insights into the specific transcriptional program operated by a given tissue Treg cell population in limiting ongoing local inflammation, we used scatter plots to compare genes that were differentially expressed in Treg cells *vs.* Tconv cells isolated from a particular tissue in mice with or without DT treatment (**Fig. 1D-F**). To this end, only genes that are upregulated (red) or downregulated (blue) in Treg cells when compared to Tconv cells under DT-treated condition by more than 1.5 fold and at least 1.5 fold higher or lower than those under PBS-treated control condition, respectively, were considered to be potentially involved in Treg cell-mediated control of ongoing systemic autoimmunity. Moreover, through Venn diagrams, we further identified genes that are commonly regulated in all or several Treg cell populations *vs.* ones that are uniquely up- or down-regulated in a given Treg cell subset over their Tconv cell counterpart from a particular anatomical location during autoimmunity (**Fig. 1G**). Finally, Gene ontology (GO) term enrichment analysis of differentially expressed genes in Treg cells isolated from different tissues under DT-treated condition revealed biological pathways that could be potentially critical for tissue Treg cell-mediated control of autoimmune inflammation in a given tissue microenvironment (**Fig. 1H** and **Fig. S2**). Among the identified biological processes, genes related to leukocyte proliferation could be observed in all Treg cell populations. On the other hand, genes related to regulation of T cell activation were specifically enriched in the intestinal Treg cells (**Fig. 1H**). These resulted suggested that while transferred Treg cells were all undergoing rapid expansion to control ongoing autoimmunity, regulation of T cell activation in the gut-associated tissue required a specialized suppressor program employed by the intestinal Treg cells.

### IL-27 is specifically induced in gut-associated Treg cells isolated from inflammatory conditions

As discussed above, despite the accumulation of knowledge of tissue Treg cells over the past decade, the precise mechanisms by which those tissue Treg cells control their corresponding local immune responses remains poorly understood. To directly address this important issue, we further explored the identified tissue Treg cell gene signatures with a focus on the ones encoding for effector molecules that have been previously associated with Treg cell suppressor program or a role in immune regulation (**Suppl. Table I** and **II**). Among the genes identified in the Venn diagram analysis that are selectively upregulated in intestinal Treg cells from mice with ongoing autoimmune-mediated inflammation (**Fig. 1G**, top), *Il27,* which encodes a subunit (IL-27p28) of a heterodimeric cytokine, IL-27, is of particular interest (**Fig. 2A**). IL-27, composed of EBI3 and IL-27p28, is a pleiotropic cytokine of the IL-6/IL-12 superfamily. While the initial studies pointed to IL-27 as a cytokine that can promote Th1 immunity, subsequent research revealed a more robust immunoregulatory role of IL-27 in controlling a diverse range of immune responses in many different diseases ^23^. IL-27 can directly inhibit the production of IL-17 and GM-CSF ^28–31^. Moreover, IL-27 has also been shown to exert its suppressive effects indirectly through inducing IL-10 production by many T cell subsets or via promoting a specific T-bet^+^Foxp3^+^Treg cell subset specialized to limit Th1 immunity ^32–34^. Nevertheless, even though the immunomodulatory role of IL-27 has now been clearly recognized, the impact of IL-27 derived from different cellular sources on immune regulation appears to be very complex ^35^. Moreover, even though Treg cells are known to express high levels of EBI3 ^36^, a finding that is also supported by our RNA-seq studies (**Fig. 2B**), it was IL-35, another heterodimeric cytokine composed of EBI3 and IL-12α rather than IL-27p28, that has been previously shown to serve as a major Treg cell suppressor molecule ^36^. However, unlike *Il27* and *Ebi3*, the level of *Il12a* expression in Treg cells remained low and unaltered irrespective of tissue origin or inflammatory condition (**Fig. 2C**). These findings from the RNA-seq study were confirmed by qPCR analysis (**Fig. 2D-F**). Moreover, by taking an ELISA approach in which IL-27p28 and EBI3 (for IL-27) or IL-12α and EBI3 (for IL-35) were captured and detected, respectively, we have also selectively detected a significantly increased amount of IL-27 protein but not IL-35 protein produced by intestinal Treg cells from mice with DT treatment (**Fig. 2G** and **H**).

**Figure 2.**
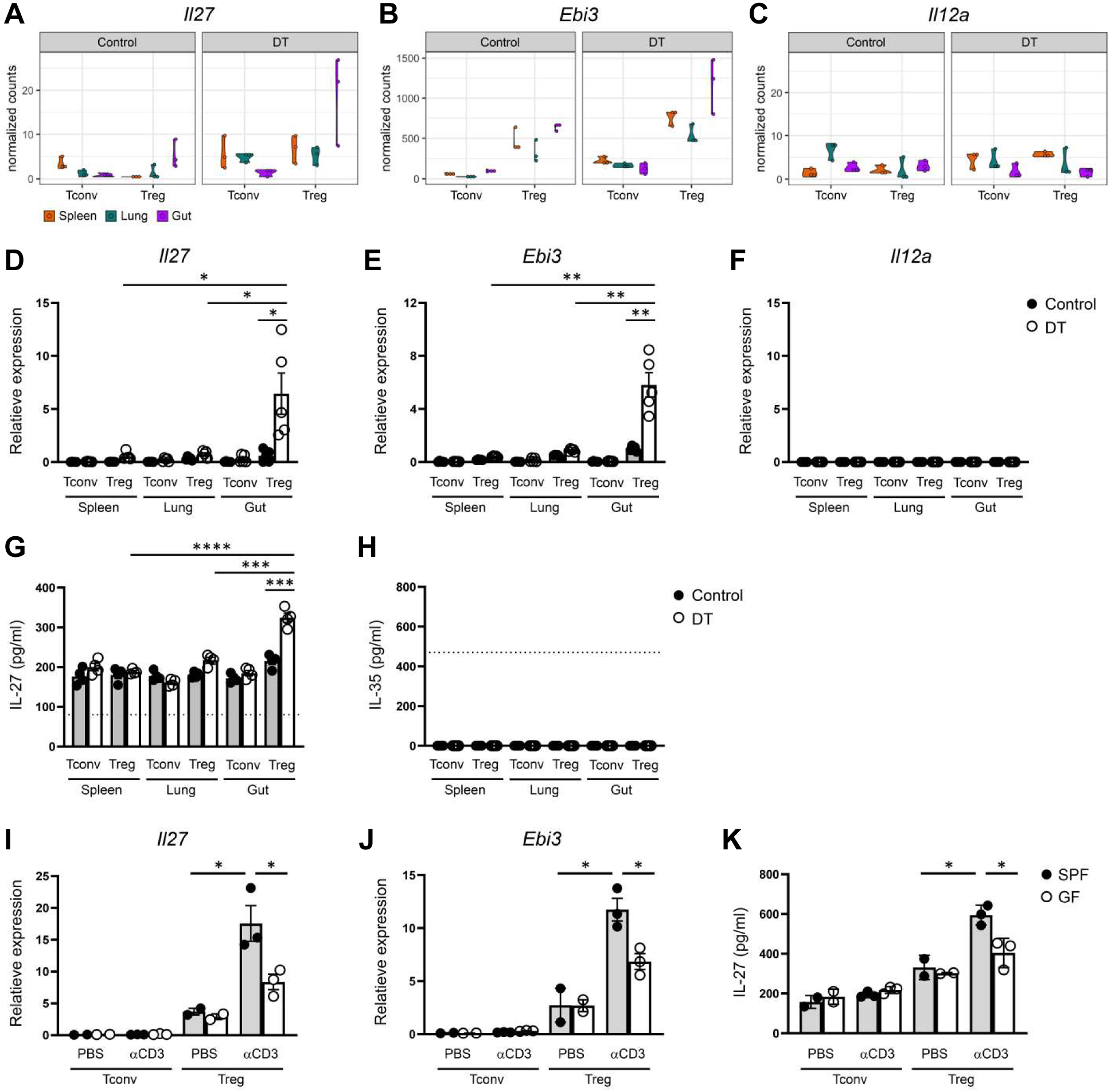
Identification of IL-27 not IL-35 as a potential suppressor molecule specifically produced by intestinal Treg cells under autoimmune inflammation. Violin plots of **(A)** *Il27*, **(B)** *Ebi3*, and **(C)** *Il12a* in Tconv and Treg cells in different tissues from control PBS- or DT-treated *Foxp3^DTR^* mice from RNA-seq analysis. qPCR analyses for the expressions of **(D)** *Il27*, **(E)** *Ebi3*, and **(F)** *Il12a* in Tconv and Treg cells in different tissues from control PBS- or DT-treated mice. ELISA analyses of the production of **(G)** IL-27 or **(H)** IL-35 by Tconv and Treg cells in different tissues from control PBS- or DT-treated mice. qPCR analyses for the expressions of **(I)** *Il27* and **(J)** *Ebi3* in Tconv and Treg cells in SI from control PBS- or aCD3-treated SPF or GF mice. ELISA analyses of the production of **(K)** IL-27 by Tconv and Treg cells in SI LP from control PBS- or aCD3 mAb-treated SPF and GF mice. Dotted line represents the minimum detection limit of the indicated cytokine. Each symbol represents an individual mouse, and the bar represents the mean. Data are pooled from at least two independent experiments. *, P < 0.05; **, P < 0.01.

Next, we sought to determine whether elevated expression of IL-27 in intestinal Treg cells could also be observed in a different inflammatory setting other than the aforementioned condition in which severe autoimmunity was induced by global Treg cell ablation. To this end, we employed an anti-CD3 (αCD3) monoclonal antibody (mAb)-induced intestinal disease model in which treatment of αCD3 mAb has been shown to lead to acute inflammation and intestinal pathology primarily in the small intestine ^37^. As shown in **Fig. 2I-K**, significant increased expressions of both transcript and protein of IL-27 were also detected in gut Treg cells in mice receiving αCD3 mAb treatment. These results suggested that elevated secretion of IL-27 by intestinal Treg cells is likely a common phenotype that could be detected in various inflammatory settings. Interestingly, such increases in IL-27 expression were greatly diminished in germ-free (GF) mice when compared to specific-pathogen-free (SPF) controls, implying that in addition to inflammation, commensal bacteria play a major role in the induction of IL-27 in intestinal Treg cells (**Fig. 2I-K**). It should be noted that elevated production of IL-10, a known Treg cell suppressor molecule, by gut Treg cells was also detected in our analysis (**Fig. S3**). However, unlike IL-27, high levels of IL-10 expression in gut Treg cells could already be observed at steady state with no clear further upregulation even in the presence of autoimmune inflammation. Given that gut-associated Treg cells selectively upregulated IL-27 expression in mice with systemic immunity, our results suggested that while Treg cell-derived IL-10 has been long recognized for its role in maintaining gut homeostasis ^18^, IL-27 produced by Treg cells likely plays a more active role in controlling ongoing intestinal inflammation.

### A selective increase in Th17 responses in the intestine of mice with Treg cell-specific IL- 27 ablation

To directly examine the function of IL-27 in Treg cell-mediated immune regulation particularly in the gut-associated tissue, we generated mice with Treg cell-specific deletion of IL-27p28 (*Foxp3^Cre^Il27^fl/fl^*; Treg-KO). Consistent with the hypothesized role of IL-27, or the lack thereof, in general Treg cell biology, Treg-KO mice did not develop any obvious immune phenotype or autoimmune pathology even when they aged (more than 6 months of age; **Fig. 3**, **Fig. S4** and data not shown). The frequencies and numbers of Treg cells in both lymphoid and non-lymphoid tissues including both small and large intestine were comparable between Treg-KO mice and their control littermates (**Fig. 3A** and **Fig. S4A, B**). The suppression capacity of Treg cells isolated from either the intestinal tissue or the spleen based on the *in vitro* suppression assay was also not impeded by the absence of IL-27 production (**Fig. 3B, C** and **Fig. S4C**). Consequently, Tconv cells remained under control as no difference in their proliferation and activation in Treg-KO mice compared to their control littermates could be observed (**Fig. 3D, E** and **Fig. S4A, B**). Interestingly, despite exhibiting no detectable inflammatory responses, the loss of IL-27p28 in Treg cells already resulted in a selective increase in the production of IL-17 by Teff cells in the gut-associated tissue from mice at the age of 8-12 weeks even in the absence of any immunological challenges (**Fig. 3F, G** and **Fig. S4A**). Moreover, supporting our proposed Treg cell-intrinsic role of IL-27 in the intestine, the dysregulated IL-17 phenotype in mice with Treg cell-specific IL-27p28 ablation was only observed in the intestine as no alteration in Th17 responses was found in lymphoid and other non-lymphoid tissues (**Fig. S4B** and data not shown). It should be noted, however, that dendritic cells (DCs) and other myeloid cells as well as intestinal epithelial cells (IECs) have all been recently shown to regulate intestinal homeostasis through the production of IL-27 ^35^. Considering the reported role of IL-27 in directly inhibiting Th17 differentiation ^29^, it is possible that IL-27 produced by different gut-resident cells could all contribute to the regulation of intestinal Th17 immunity. To test this possibility, we examined Th17 responses in mice with DC- (DC-KO), myeloid cell- (Mye-KO) and IEC-specific (IEC-KO) deletion of IL-27p28, respectively. As shown in **Fig. S5A-C**, we did not observe any alteration in Th17 cell frequencies in the aforementioned mouse lines compared to their corresponding littermate controls. In contrast, in mice harboring T cells incapable of responding to IL-27 (T-rKO), elevated Th17 responses similar to those found in Treg-KO mice were easily detected (**Fig. S5D** and **E**). Finally, to further confirm that the observed Th17 phenotype in Treg-KO mice is indeed due to the specific loss of Treg cell-derived IL-27 rather than the potential off-target deletion driven by Foxp3-Cre as reported previously ^38, 39^, we performed the adoptive transfer study in which IL-27-deficient Treg cells were co-transferred with congenically marked Foxp3^-^CD4^+^ T cells isolated from *Foxp3^KO^* mice into RAG-deficient mice. Previously, it has been shown that elevated Th1, Th2 and Th17 responses in the Foxp3-deficient T cell compartment could be controlled upon co-transfer of Treg cells ^40^. To this end, consistent with what we found in Treg-KO mice, IL-27-deficient Treg cells also selectively failed to restrain Th17 responses in the gut in the adoptive transfer study in which only Treg cells are incapable of producing IL-27 (**Fig. 3H, I** and **Fig. S4D**). Collectively, these results indicate that Treg cell-derived IL-27, despite being produced at a low level, plays a non-redundant role in fine-tuning Th17 responses particularly in gut-associated tissue. Our data also support previous observations of distinct roles of IL-27 produced by different cell types in maintaining intestinal homeostasis ^35^.

**Figure 3.**
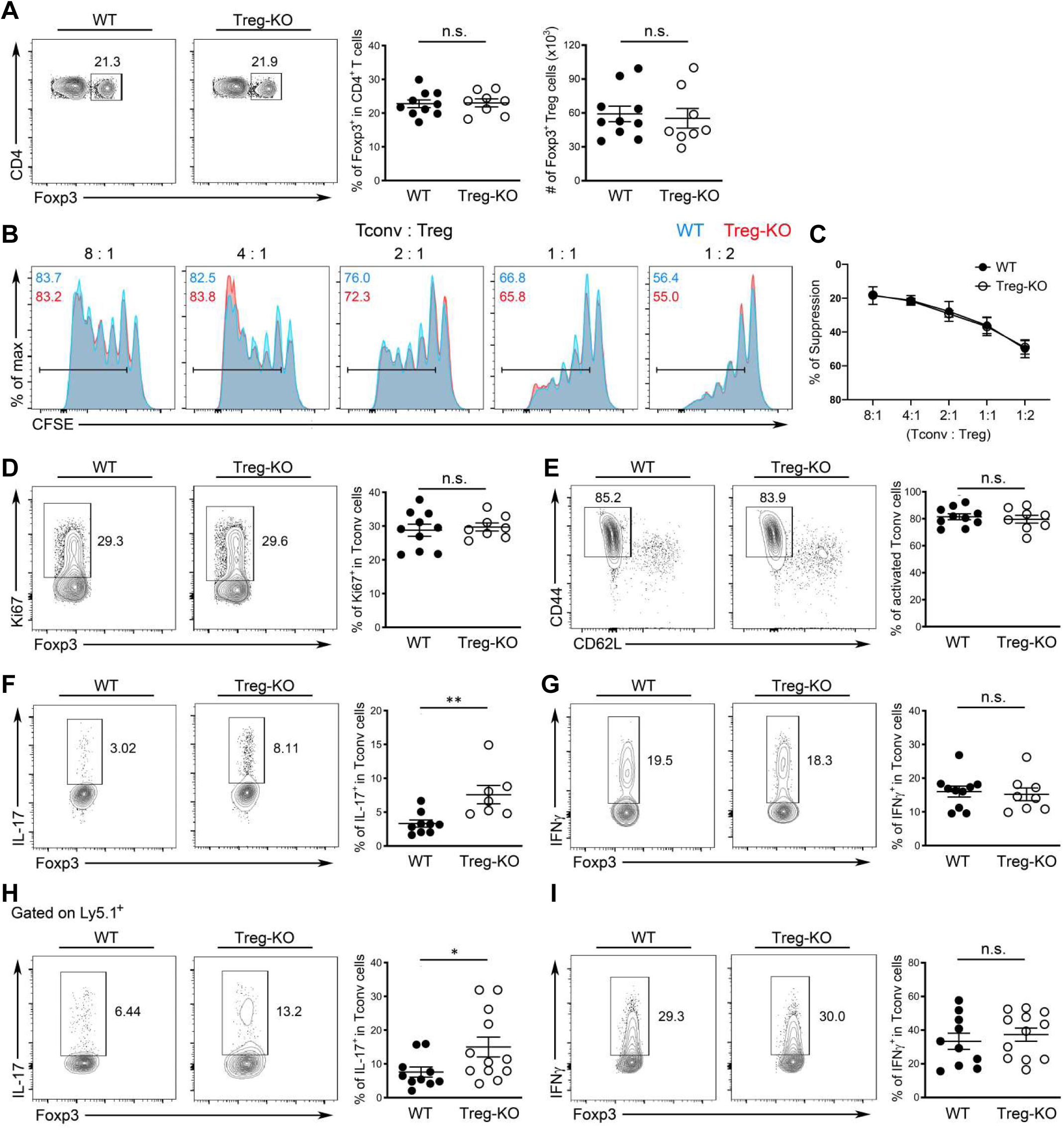
A selective defect in regulating intestinal Th17 responses in mice harboring Treg cells incapable of producing IL-27. **(A)** FACS analysis, frequencies and numbers of Foxp3^+^ Treg cells in SI LP of Treg-KO mice and WT littermates (∼8-12 weeks). **(B)** FACS analysis and **(C)** % of suppression of proliferation of WT Tconv cells by SI LP Treg cells isolated from either WT or Treg-KO mice in an *in vitro* suppression assay. Data are representative of three independent experiments. n = 6. FACS analysis and frequencies of **(D)** Ki67^+^, **(E)** CD44^hi^CD62L^lo^, **(F)** IL-17^+^, and **(G)** IFNg^+^ Tconv cells in SI LP of Treg-KO mice and WT littermates (∼8-12 weeks). FACS analysis and frequencies of **(H)** IL-17^+^ and **(I)** IFNg^+^ Ly5.1^+^ Tconv cells in SI LP of RAG-deficient mice three weeks after co-transferred with Treg cells isolated from either Treg-KO mice or WT littermates. Each symbol represents an individual mouse, and the bar represents the mean. Data are pooled from at least three independent experiments. n.s., not significant; *, P < 0.05; **, P < 0.01.

### Loss of Treg cell-derived IL-27 led to exacerbated intestinal inflammation and colitis-associated tumor

Although we found a role for Treg cell-derived IL-27 at steady state, since *Il27* is upregulated in intestinal Treg cells during autoimmunity, we hypothesized that deletion of IL-27 in Treg cells would lead to an even stronger IL-17 response and cause more severe intestinal immunopathology in mice with ongoing inflammation. To this end, we went back to the aforementioned αCD3 mAb-driven intestinal disease model in which pro-inflammatory Th17 cells were shown to be predominantly responsible for the development of intestinal pathology ^41, 42^. To examine the functional impact of IL-27 ablation in Treg cells on regulating intestinal inflammation, we followed the mice challenged with αCD3 mAb over time and euthanized the mice one day after the last αCD3 mAb injection for FACS and histopathology analysis. In agreement with our previous studies, we could detect significantly increased IL-27, but not IL-35, secretion by gut Treg cells isolated from αCD3 mAb treated mice and the production of IL-27 was completely abolished in Treg-KO mice (**Fig. S6A** and **B**). Moreover, as shown in **Fig. 4A** and **B**, we found that Treg-KO mice exhibited more pronounced inflammation-induced weight loss along with more severe gut pathology compared to their control littermates. The exacerbated disease phenotype was not due to insufficient Treg cell numbers as Treg cells from both Treg-KO mice and their controls were able to expand to a similar degree in the attempt to control the αCD3 mAb-induced intestinal inflammation (**Fig. 4C**). Nevertheless, consistent with what we observed during homeostasis, Treg cells incapable of producing IL-27 exhibited a selective defect in regulating Th17 cells as markedly increased IL-17 but not IFNγ responses were observed in Treg-KO mice compared to WT littermates (**Fig. 4D** and **E**). Similar results were also obtained from mice harboring T cells unresponsive to IL-27 while comparable Th17 responses were observed between mice with DC-, myeloid cell- and IEC-specific deletion of IL-27 and their corresponding WT controls upon αCD3 mAb administration (**Fig. S5F-J**), indicating that Treg cells remain the major cellular source of IL-27 required to control intestinal Th17 responses under inflammatory conditions.

**Figure 4.**
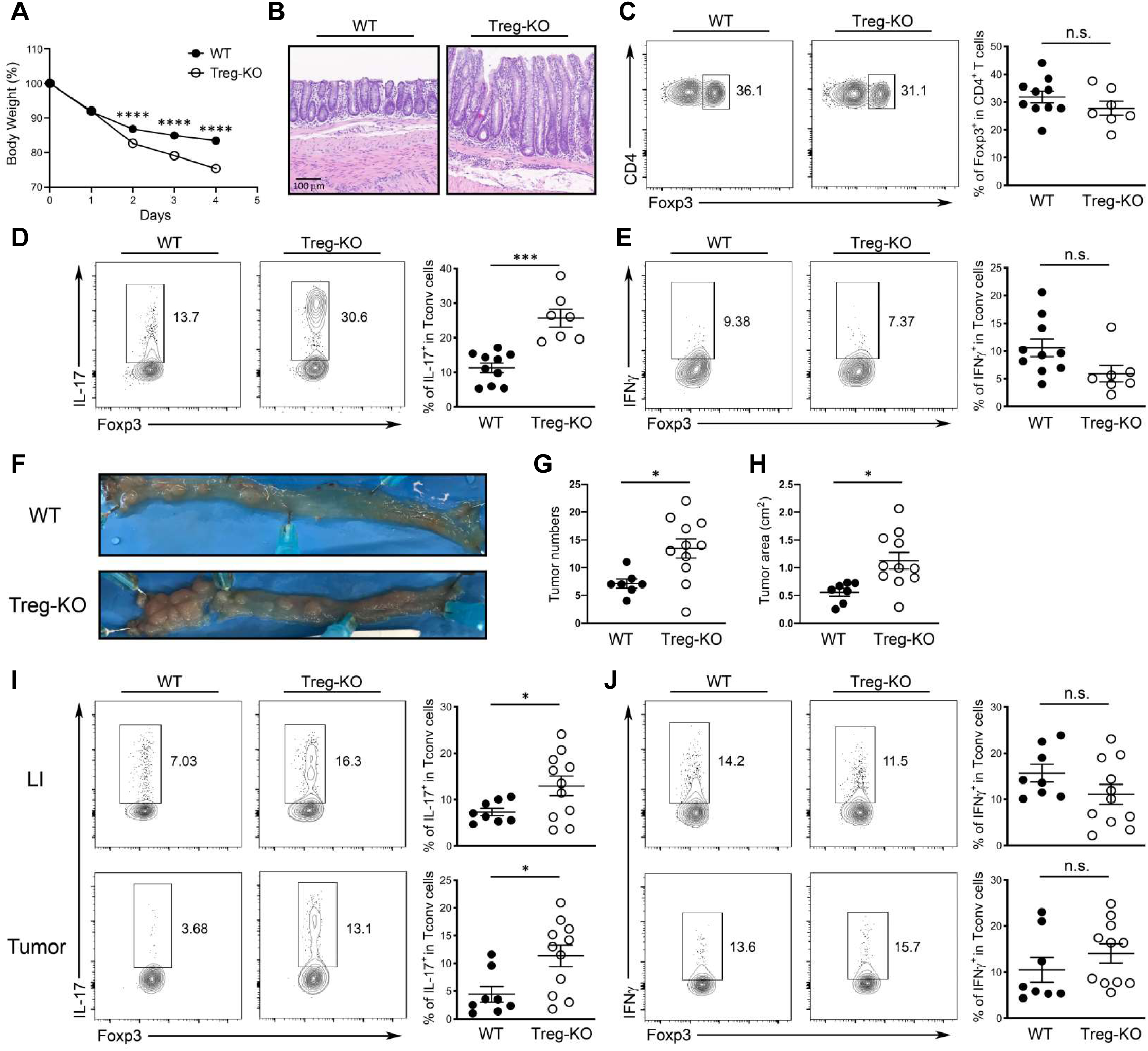
Loss of Treg cell-derived IL-27 led to exacerbated intestinal inflammation and colitis-associated cancer. **(A)** Percentage of body weight change of Treg-KO mice and WT littermates after aCD3 mAb administrations. **(B)** 4 days after initial aCD3 mAb injection, small intestine sections from the mice were stained with H&E for microscopic imaging. Scale bar: 100 μm. FACS analysis and frequencies of **(C)** Foxp3^+^ Treg cells and **(D)** IL-17^+^, and **(E)** IFNg^+^ Tconv cells in SI LP of aCD3 mAb-treated Treg-KO mice and WT littermates. **(F)** Representative colonic photos, **(G)** numbers, and **(H)** area of colorectal tumors in Treg-KO mice and WT littermates 12 weeks after AOM/DSS treatment. FACS analysis and frequencies of **(I)** IL-17^+^ and **(J)** IFNg^+^ Tconv cells in LI LP or colorectal tumors of Treg-KO mice and WT littermates 12 weeks after AOM/DSS treatment. Each symbol represents an individual mouse, and the bar represents the mean. Data are pooled from at least three independent experiments. n.s., not significant; *, P < 0.05; ***, P < 0.001; ****, P < 0.0001.

In addition to αCD3 mAb-induced intestinal inflammation, elevated Th17 cell responses were also previously shown to promote tumorigenesis in an Azoxymethane (AOM)/Dextran sulfate sodium (DSS) model of carcinogen-induced colitis-associated cancer (CAC) ^43, 44^. Interestingly, even though the role of Treg cells in controlling anti-tumor immunity has been well established, during CAC tumorigenesis, many studies have suggested that Treg cells could exhibit either anti- or pro-tumor function depending upon the timing during tumor development or progression ^45^. To this end, transient deletion of Treg cells during early phase of CAC has been shown to exacerbate intestinal inflammation that could promote tumorigenesis. It is thus plausible that loss of Treg cell-derived IL-27-mediated regulation of Th17 responses in the intestine could create a microenvironment favorable for tumor growth. Indeed, supporting our proposed role of Treg cell-derived IL-27 in limiting Th17-driven tumorigenesis, as shown in **Fig. 4F-H**, increased colon tumor burdens were observed in Treg-KO mice compared to WT littermates three months after the AOM and DSS treatment. Consistent with our observations in the αCD3 mAb-induced intestinal disease study, selective increases in the frequencies of Th17 cells but not IFNγ-producing Th1 cells could be found in both large intestine (LI) and tumors in mice harboring Treg cells incapable of producing IL-27 (**Fig. 4I** and **J**). Together, by using both acute and chronic intestinal inflammatory disease models, our results demonstrate a critical role of Treg cell-derived IL-27 in limiting Th17-driven gut pathology.

### Treg cell-derived IL-27 is dispensable to control Th17-driven experimental autoimmune encephalomyelitis

Thus far, our studies have shown that Treg cell-specific IL-27 ablation resulted in a selective defect in the regulation of Th17 responses in the intestinal tissue. However, while Treg cell-derived IL-27 does not seem to play a noticeable role in tissues other than the gut at steady state, it remains plausible that IL-27 produced by Treg cells might still be required to control Th17 immunity outside of the intestinal tissues when Th17-driven inflammatory responses are triggered. To test this possibility, we used a model of experimental autoimmune encephalomyelitis (EAE), a central nervous system (CNS) autoimmune disorder in which autoreactive Th1 cells and Th17 cells have both been suggested to serve as central mediators in promoting disease pathogenesis ^46^. Moreover, during EAE, IL-27 has also been previously reported to limit neuroinflammation through directly suppressing the development of Th17 cells ^28^. Consistent with previous studies, upon EAE induction, we detected elevated Th17 responses in the CNS of mice harboring T cells incapable of responding to IL-27 whereas the frequencies of IFNγ-secreting cells remained unchanged (**Fig. 5A** and **B**). As a result, compared to their WT littermates, a significant worsening of EAE was found in mice harboring T cells incapable of responding to IL-27 (**Fig. 5C**). However, in contrast to our findings with the intestinal inflammation models, no alteration in the frequencies of Th17 (and Th1) cells could be found in Treg-KO mice upon EAE induction (**Fig. 5D** and **E**). Consequently, both Treg-KO mice and their WT littermates exhibited similar disease phenotypes (**Fig. 5F**). Altogether, while our results have confirmed a T cell-intrinsic role of IL-27 signaling in restricting Th17 responses, unlike what was observed in the intestine, Treg cell-derived IL-27 does not play a functional role in the CNS even in the presence of Th17-driven neuroinflammation.

**Figure 5.**
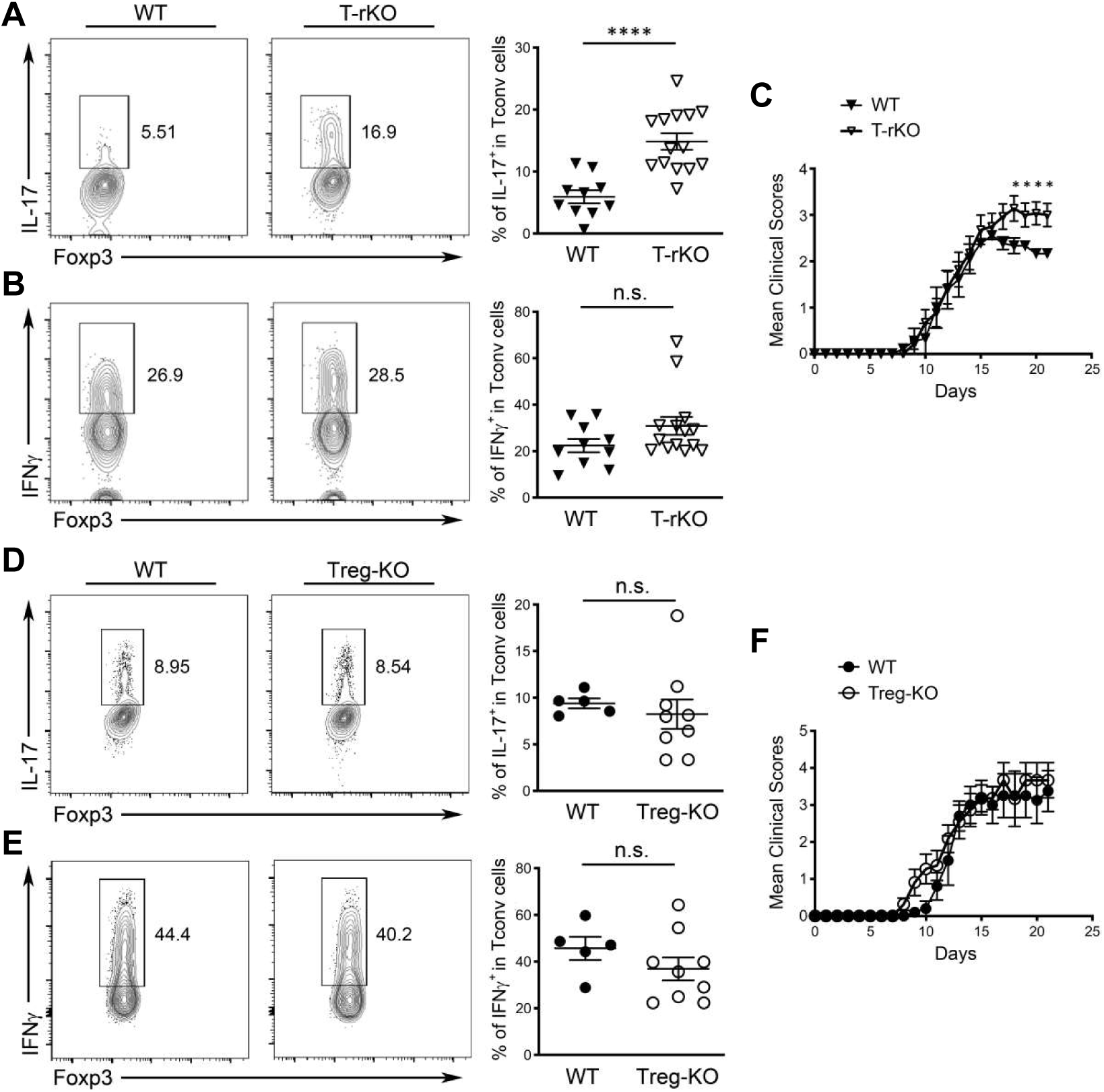
Treg cell-derived IL-27 is dispensable to control Th17-driven experimental autoimmune encephalomyelitis. FACS analysis and frequencies of **(A)** IL-17^+^ and **(B)** IFNg^+^ Tconv cells in the brain of T-rKO mice and WT littermates 21 days after EAE induction. **(C)** The disease severity was scored regularly based upon clinical symptoms. FACS analysis and frequencies of **(D)** IL-17^+^ and **(E)** IFNg^+^ Tconv cells in the brain of Treg-KO mice and WT littermates 21 days after EAE induction. **(F)** The disease severity was scored regularly based upon clinical symptoms. Each symbol represents an individual mouse, and the bar represents the mean. Data are pooled from at least two independent experiments. n.s., not significant; *, P < 0.05; ****, P < 0.0001.

### Enhanced IL-17 responses in mice with Treg cell-specific IL-27 ablation helped protect against enteric bacterial infection

While uncontrolled Th17 responses have been frequently associated with intestinal immunopathology, IL-17, the main effector cytokine produced by Th17 cells is crucial for protecting against extracellular bacteria, fungi and viruses at the mucosal surface ^47^. Previously, it was shown that *Citrobacter rodentium* (*C. rodentium*), a naturally occurring mouse pathogen that preferentially impacts the colon, can induce a strong local Th17 response that is necessary for protection ^48^. It is thus possible that the enhanced IL-17 responses triggered by the absence of Treg cell-derived IL-27 could help the host control infection with enteric pathogens such as *C. rodentium*. To this end, similar to what we found in the aforementioned autoimmune-driven inflammation models, we could also detect substantial amount of IL-27 but not IL-35 secreted by Treg cells isolated from the colon of mice that were infected with *C. rodentium* whereas only minimal IL-27 production by Treg cells isolated from the spleen in the same infected mice could be observed (**Fig. S6C** and **D**). It should be noted that in agreement with a previous report ^49^, during infection, Tconv cells were also capable of producing IL-27 as shown in our analysis (**Fig. S6C** and **D**). Nevertheless, the amount of IL-27 secreted by intestinal Treg cells was still much higher compared to what could be found by Tconv cells (**Fig. S6D**). Finally, like what we found in Treg cells from the αCD3 mAb-induced intestinal disease model, these results confirmed the previous notion that elevated expression of IL-27 by gut-associated Treg cells is not just due to the non-physiological nature of the severe autoimmunity caused by global Treg cell ablation. Inflammation resulting from either local autoimmune responses or natural infections in the intestine could similarly drive IL-27 induction in Treg cells.

Next, we sought to determine the effect of Treg cell-specific IL-27 ablation on host defense against this enteric pathogen. As shown in **Fig. 6A** and **B**, we found that *C. rodentium*-infected Treg-KO mice exhibited further elevated frequencies of IL-17^+^ but not IFNγ^+^ Teff cells in the colon compared to their WT littermates. Consequently, significantly reduced bacterial burden in Treg-KO mice over littermate controls was observed (**Fig. 6C**). These results support our hypothesis that Treg cell-derived IL-27 regulates intestinal Th17 immunity. Moreover, it is also interesting to speculate that inducing IL-27 secretion by gut Treg cells might be a strategy utilized by Th17-driven enteric pathogens such as *C. rodentium* to evade the host’s immune system. On the other hand, it is uncertain as to whether IL-27 produced by Treg cells would have a similar impact on the host defense against other pathogens in the intestine when a different type of immune responses is triggered. To address this issue, we employed a *Toxoplasma gondii* (*T. gondii*) infection model in which a strong IFNγ-mediated Th1 response is induced in the gut necessary for the clearance of this enteric pathogen ^50^. As shown in **Fig. 6D** and **E**, while increased frequencies of IL-17-producing Th17 cells could still be observed in Treg-KO mice during *T. gondii* infection, comparable Th1 responses were elicited in Treg-KO mice and their WT littermates. Moreover, both Treg-KO mice and littermate controls were able to control pathogen efficiently (**Fig. 6F**). Collectively, our results from two different enteric infection models further demonstrate a selective regulatory effect of Treg cell-derived IL-27 on Th17 immunity, but not Th1 immunity, during intestinal infections.

**Figure 6.**
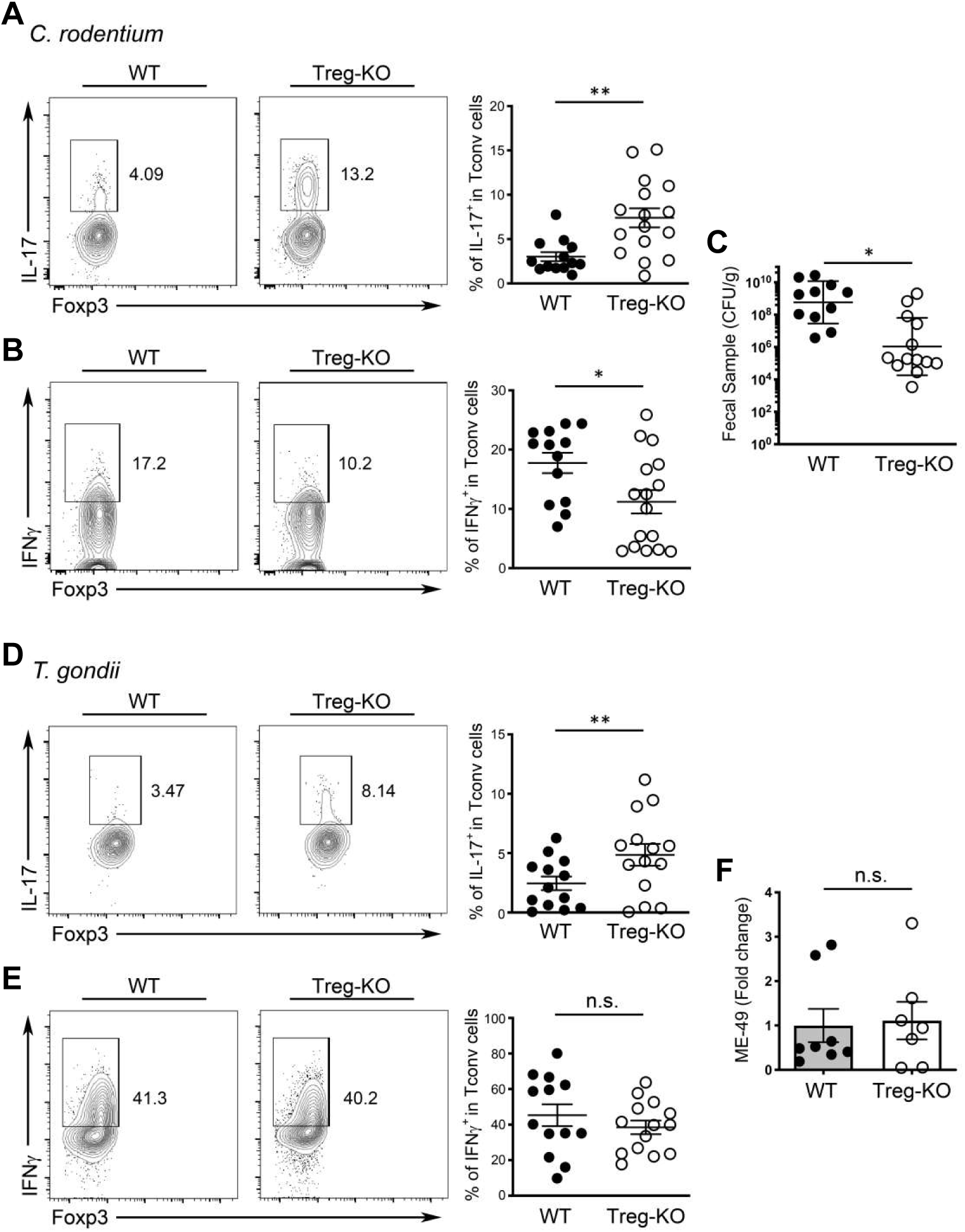
Enhanced IL-17 responses in mice with Treg cell-specific IL-27 ablation selectively helped protect against enteric bacterial infection. FACS analysis and frequencies of **(A)** IL-17^+^ and **(B)** IFNg^+^ Tconv cells in LI LP of Treg-KO mice and WT littermates at day 10 post *C. rodentium* infection. **(C)** Enumeration of *C. rodentium* in the feces of Treg-KO mice and WT littermates at day 10 post *C. rodentium* infection. FACS analysis and frequencies of **(D)** IL-17^+^ and (E) IFNg^+^ Tconv cells in SI LP of Treg-KO mice and WT littermates at day 8 post *T.gondii* infection. **(F)** qPCR analysis of parasite burden in small intestine of Treg-KO mice and WT littermates at day 8 post *T.gondii* infection. Each symbol represents an individual mouse, and the bar represents the mean. Data are pooled from at least three independent experiments. n.s., not significant; *, P < 0.05; **, P < 0.01.

### Single cell-transcriptomic analysis revealed a distinct intestinal Treg cell subset responsible for the production of IL-27 particularly under inflammatory conditions

Thus far, our studies have demonstrated that intestinal Treg cells are capable of secreting IL-27 to control Th17 responses particularly under inflammation. However, as several intestinal Treg cell populations have been reported to maintain gut homeostasis and intestinal tolerance ^51–54^, it remains unclear as to whether expression of IL-27 is a common feature for the entire intestinal Treg cell population or there is a specific Treg cell subset primarily responsible for IL-27 production. To further characterize IL-27-expressing Treg cells, we performed single-cell RNA sequencing (scRNA-seq) analysis of intestinal Treg cells from mice with or without intestinal inflammation caused by *C. rodentium* infection. As shown in **Fig. 7A**, by using dimensional reduction by Uniform Manifold Approximation and Projection (UMAP) for visualization of intestinal Treg cells, we observed 4 distinct Treg cell clusters (R0-R3). Treg cells from R0 cluster exhibits high expression of *Tcf7* but not *Sell* resembling the activated Treg cells that was previously reported (**Fig. 7B**) ^55^. On the other hand, R1 cluster exhibits high expression of genes resembling effector Treg cells (e.g. *Il10* and *Gzmb*) while R2 cluster expresses transcripts (e.g. *Sell* and *Bach2*) indicative of resting Treg cells (**Fig. 7B** and **Fig. S7A** and **B**) ^56, 57^. Finally, R3 cluster is enriched with *Il1rl1* (i.e. *St2*) expressing cells (**Fig. 7B** and **Fig. S7C**). Interestingly, while the distribution of these 4 clusters was rather comparable between control and Citrobacter-infected groups (**Fig. 7C**), *Il27* transcripts could only be detected in the R0 cluster from the infected group (**Fig. 7D**). Unfortunately, as reported previously ^58^, due to the inherent lack of sensitivity of scRNA-seq for low-abundance transcripts such as cytokines, only few *Il27* transcript signals were detected. Nevertheless, we do not think this is merely an experimental artifact as the signals of *Ebi3*, a gene encoding another subunit of IL-27 that has been well-recognized to be highly expressed in Treg cells were also barely detectable (**Fig. 7E**). Unlike *Il27*, signals of *Ebi3* could be found in both R0 and R1 clusters, suggesting that R0 cluster is enriched with IL-27 expressing Treg cells while R1 cluster likely contains Treg cells that could produce other cytokines consisting of EBI3.

**Figure 7.**
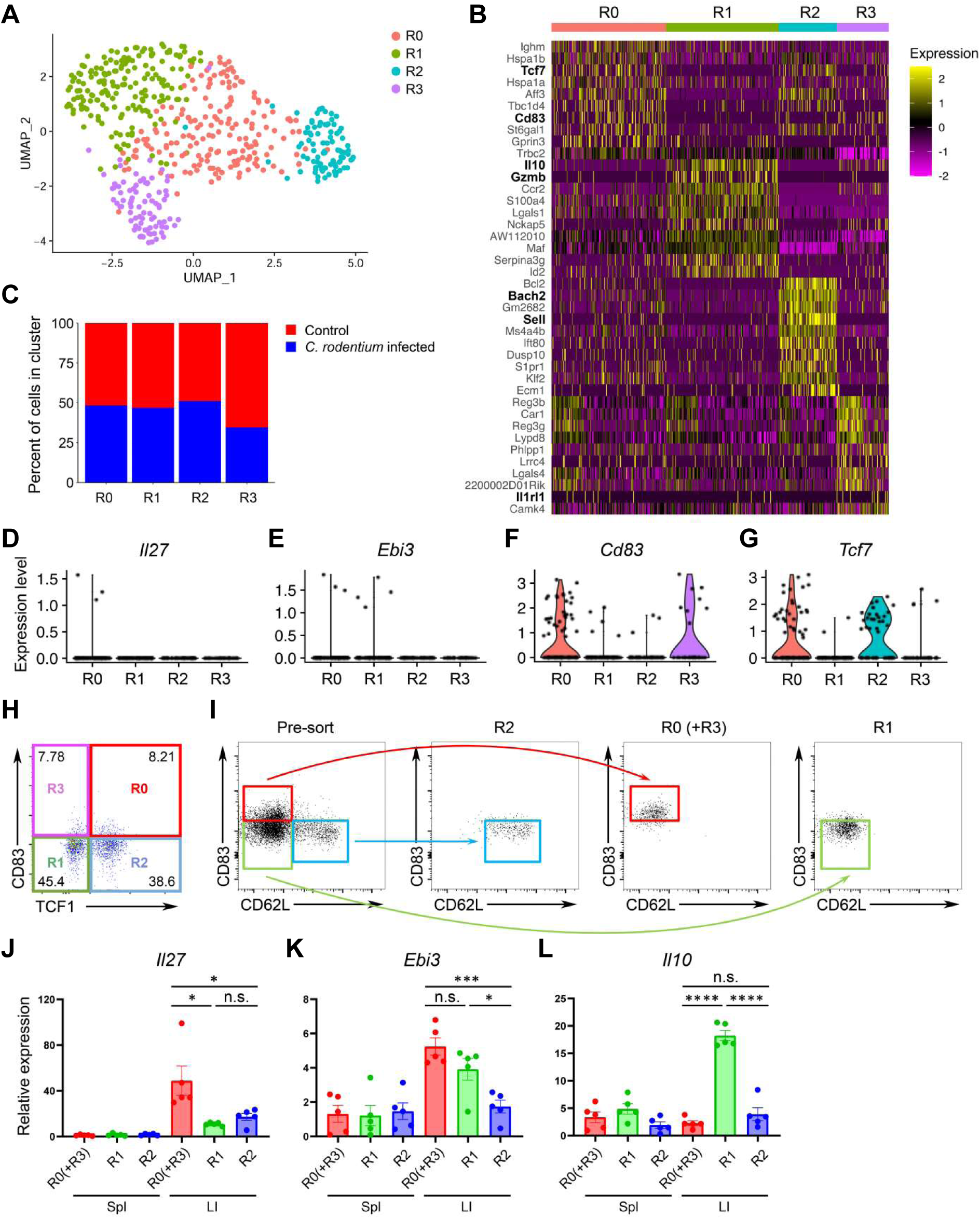
Single-cell analyses identify a putative IL-27-producing Treg cell subset in the inflammatory intestinal tissue. **(A)** UMAP plots of intestinal Treg cell clusters, colored by cluster identity. **(B)** Heatmap of top 10 differently expressed genes between each intestinal Treg cell clusters. **(C)** Percentage of cells within each intestinal Treg cell cluster from mice with or without *C. rodentium* infection. Violin plots of **(D)** *Il27*, **(E)** *Ebi3*, **(F)** *Cd83*, and **(G)** *Tcf7* in different intestinal Treg cell clusters from *C. rodentium-* infected mice. **(H)** FACS analysis of intestinal Treg cell clusters based on the expression fo CD83 and TCF1. **(I)** Representative of FACS profiles with gating strategy for isolating different intestinal Treg cell subsets. qPCR analyses for the expressions of **(J)** *Il27*, **(K)** *Ebi3*, and **(L)** *Il10* in different Treg cell subsets in spleen or LI LP from *C. rodentium-* infected mice. Each symbol represents an individual mouse, and the bar represents the mean. Data are pooled from two independent experiments. n.s., not significant; *, P < 0.05; ***, P < 0.001; ****, P < 0.0001.

To further confirm our scRNA-seq results, we first FACS analyzed intestinal Treg subsets from the infected samples based on the expressions of CD83 and TCF1. CD83, a heavily glycosylated Ig-like type 1 transmembrane protein which has been previously shown to be a marker for activated Treg cell lineages ^59^, was found to be one of top 10 highly expressed genes along with *Tcf7* (the gene encoding TCF1) in R0 cluster (**Fig. 7B**). Moreover, in addition to R01 cluster, *Cd83* and *Tcf7* could also be detected in R3 and R2 cluster to a lesser degree, respectively (**Fig. 7F** and **G**). Thus, by utilizing the expression patterns of these two molecules, we could divide intestinal Treg cells into 4 clusters identified by our scRNA-seq analysis: R0: CD83^+^TCF1^+^; R1: CD83^-^TCF1^-^; R2: CD83^-^TCF1^+^; R3: CD83^+^TCF1^-^ (**Fig. 7H**). Next, without having the TCF1 reporter mice in hands, we then used CD62L (encoded by *Sell*) as a surrogate for TCF1 as it is also highly expressed in R2 cluster (**Fig. 7B** and **Fig. S7B**). While we could not easily separate R0 and R3 clusters since both clusters are enriched with *Cd83*-but not *Sell*-expressing cells, three populations of Treg cells: R0+R3 (CD83^+^CD62L^lo^; Red), R1 (CD83^-^ CD62L^lo^; Green), and R2 (CD83^-^CD62L^hi^; Blue) clusters, based on the expression patterns of CD83 and CD62L were FACS isolated (**Fig. 7I**). As shown in **Fig. 7J** and **K**, R0 (and R3) cluster-enriched Treg cells expressed high levels of *Il27* and *Ebi3* while R1 cluster-enriched Treg cells only expressed *Ebi3*. On the other hand, only R1 cluster-enriched Treg cells expressed high levels of *Il10* while none of these three genes were found to be highly expressed in R2 cluster-enriched Treg cells consistent with the aforementioned effector and resting Treg cell features in R1 and R2 clusters, respectively as suggested by our scRNA-seq studies (**Fig. 7L**). It should be noted that even though CD83^+^ Treg cells could also be found in the spleen, expression of *Il27* could not be detected, further supporting our previous findings of selective *Il27* induction in intestinal Treg cells. Finally, unlike *Cd83* (and *Tcf7*), it seems that the previously identified intestinal Treg cell genes such as *Rorc* and *Gata3* could not be used to mark these IL-27-expressing Treg cells (**Fig. S7D**). Collectively, our studies have identified a novel CD83^+^ (and TCF1^+^) Treg cell subset, which is distinct from other known Treg cell populations previously reported in the intestine, responsible for IL-27 production and that these IL-27-expressing Treg cells play a crucial role in controlling intestinal Th17 immunity particularly under inflammatory conditions.

## Discussion

As opposed to the original concept that Treg cells provide a generic level of immune regulation, it is now well appreciated that there is a high level of heterogeneity in Treg cell populations in order to effectively control a wide range of immune responses in different tissue microenvironments. However, despite the considerable efforts that have been committed in numerous gene expression profiling studies from different Treg cell populations isolated from different tissues under various physiological and pathological conditions, the exact suppressor mechanisms by which different subsets of Treg cells control a specific type of immune response in a given anatomical location have yet to be identified. Here, we discover that IL-27, a pleiotropic cytokine that is known for its diverse immunomodulatory function, is selectively induced in gut-associated Treg cells under inflammatory conditions. Our data demonstrate that IL-27 produced by Treg cells plays a non-redundant role in limiting intestinal Th17 responses and that loss of Treg cell-derived IL-27-mediated regulation of Th17 immunity could lead to either detrimental or beneficial outcomes in a context-dependent manner. Finally, the scRNA-seq analysis of Tconv cells and Treg cells in the intestinal tissue from patients with active UC further suggests that the Treg-IL-27 regulatory axis may play an equally important role in controlling gut inflammatory responses in humans.

Our studies have clearly demonstrated that under both autoimmune and infection-driven inflammatory conditions, IL-27 was specifically induced in gut Treg cells, raising an important question as to what makes the intestine unique to drive IL-27 expression in Treg cells. To this end, the intestine contains the largest surface of contact between the body and the external environment. Within the gut lumen, over 100 trillions of commensal microbes from hundreds of species consistently interact with and are recognized by TLRs on the host cells to maintain intestinal homeostasis ^60^. As discussed earlier, it has been documented that the production of IL-27 by macrophages can be induced in a TLR/MyD88-dependent manner ^61^. Even though the expression of IL-27 has never been reported in Treg cells, selective deletion of MyD88 in Treg cells has been previously shown to lead to a specific defect in controlling IL-17 responses in the gut mucosa. As a consequence, mice harboring Treg cells devoid of MyD88 failed to establish intestinal tolerance and exhibited a more severe disease phenotype during colitis ^62^. Owing to a strong resemblance between the findings in mice with Treg cell-specific deletion of IL-27 and MyD88, it is intriguing to speculate that loss of IL-27 induction in MyD88-deficient Treg cells could be the underlying mechanism responsible for the dysregulated intestinal Th17 responses. Furthermore, dysbiotic gut microbiota, particularly under inflammatory conditions, likely serve as important environmental factors to drive the expression of IL-27 in intestinal Treg cells, a notion that is directly supported by our analysis of GF animals upon αCD3 mAb-induced intestinal inflammation. Nevertheless, future studies employing whole genomic sequencing and gnotobiotic approaches are required to identify specific microbes or minimal commensal consortium that functionally contribute to the induction of IL-27 in intestinal Treg cells.

Even though IL-27 has long been recognized for its immunoregulatory role in suppressing Th17 immunity ^28, 29^, loss of IL-27 signaling has also been shown to led to enhanced IFNγ responses ^63^. Therefore, it seems puzzling as to why we did not see an effect on IFNγ responses when IL-27 production was ablated in Treg cells. Moreover, our study suggests that only IL-27 derived from Treg cells but not from other known IL-27-producing cell populations is responsible for regulating Th17 responses in the intestine. These results were also surprising since the IL-27-expressing Treg cell subset might not even be the major subset within the intestinal Treg cells as suggested by our scRNA-seq study and the amount of IL-27 secreted by Treg cells is also not likely to be higher than that made by other major IL-27 producers in the gut. However, these findings were not completely unexpected. First, while IL-27 is capable of limiting both Th1 and Th17 responses *in vivo* ^23^, it does not seem to repress but might rather promote Th1 cell differentiation through the activation of STAT1 and the induction of T-bet ^64, 65^. On the other hand, it has been well documented that IL-27 can directly inhibit Th17 cell differentiation and function through a STAT1-dependent manner ^29^. Second, we have recently demonstrated that IL-27 secreted by DCs, other myeloid cells and IECs plays distinct roles in promoting intestinal homeostasis both at steady state and during infection ^35^. Specifically, IL-27 produced by DCs was shown to be critical for the differentiation of T-bet^+^ Treg cells, a specific Treg cell subset that is required to control IFNγ-mediated Th1 immunity ^33, 34^. These results suggested that unlike its inhibitory effect on Th17 cells, IL-27 controls Th1 cell responses indirectly through the induction of T-bet^+^ Treg cells and further implied that the IL-27-producing cells and the responding cells likely reside in close proximity and directly interact with each other to achieve such a selective effect. Supporting this notion, previously, the gut microbiota has been shown to be essential for the induction of intestinal Th17 cells ^66, 67^. Considering the aforementioned putative role of gut microbes in driving IL-27 expression in Treg cells, it is conceivable that the precise location in the intestinal tissue that the IL-27-producing Treg cells inhabit is also where the differentiation of Th17 cells takes place. It could also explain why Treg cell-derived IL-27 plays a negligible role in regulating Th17 cells in the CNS despite the fact that T cell-intrinsic IL-27 signaling is still needed to limit Th17 responses during EAE. To this end, it has been previously reported that in multiple sclerosis (MS) patients, elevated expression of IL-27 was found in astrocytes, microglia and other myeloid cells. It is thus plausible that IL-27 produced by these non-Treg CNS-resident cell types is responsible for regulating Th17-driven neuroinflammation ^68^.

Previously, a subset of GATA3^+^ST2^+^Treg cells was found to be prominent in the colon. IL-33 signaling through ST2 activates GATA3 enabling colonic Treg cells to compete in the inflammatory niche and to control Teff cell responses through maintaining optimal Foxp3 expression under intestinal inflammatory stress ^51^. In addition to GATA3^+^ST2^+^Treg cells, a substantial fraction of Treg cells in the colon and in the small intestines was reported to express RORγt, a transcription factor that is originally demonstrated to antagonize Foxp3 and to promote Th17 cell differentiation ^52–54^. These RORγt^+^Treg cells were shown to exhibit enhanced suppressive capacity and are important to maintain gut homeostasis ^52–54^. In the current study, we have further identified a distinct Treg cell subset that does not seem to resemble any previously characterized intestinal Treg cell populations. This particular CD83^+^ (and TCF1^+^) Treg cell population does not express those known effector Treg cell molecules but rather appears to be specialized in IL-27 production. Considering the complex nature of the microenvironment in the intestinal tissue, one probably should not be surprised that gut-associated Treg cells can exhibit many unique characteristics. The presence of these functional distinct Treg cell subsets in the gut implies the presence of a certain division of labor between different intestinal Treg cell subsets to coordinately maintain intestinal homeostasis; an important topic requires further investigation in the future.

In humans, several genome wide association studies have identified IL-27 as a candidate gene within a susceptibility locus for IBD, an intestinal inflammatory disorder that affects many people in the United States and worldwide ^69–71^. Significantly less IL-27 was found in people harboring the risk alleles relative to those with the non-risk alleles ^69^. These studies provided evidence linking IL-27 and IBD and suggested that the observed elevations in IL-27 in certain patients likely represent an anti-inflammatory response albeit insufficient in the attempt to control the ongoing intestinal inflammation ^72^. Nevertheless, it should also be noted that there are reports pointing to a pro-inflammatory role of IL-27 in promoting colitis ^73, 74^. These seemingly contradictory findings further demonstrated the complex nature of this cytokine as IL-27 can exert its diverse activities depending on the cell type that produces it, the cell type that responds to it as well as the location and likely the timing when the stimulation occurs. Here, our studies have clearly shown that Treg cell-derived IL-27 plays a dominant and non-redundant role in IL-27-dependent regulation of intestinal Th17 responses and should pave the way for the development of IL-27 targeting as a potential therapy in treating human gastrointestinal immune disorders. Finally, the approach taken in this study not only allowed us to identify IL-27 for the first time as a Treg cell suppressor molecule selectively required for controlling a particular type of immune response in a specific tissue location, but also established a powerful platform for future investigation into tissue-specific Treg cell suppressor programs.

## Materials and Methods

### Mice

*Foxp3^KO^* mice ^75^, *Foxp3^DTR^* mice ^27^, *Foxp3^Thy1.1^* mice ^76^, *CD11c-Cre Il27^fl/fl^* (DC-KO) mice ^77^, *Lysozyme-Cre Il27^fl/fl^* (Mye-KO) mice, *Villin1-Cre Il27^fl/fl^* (IEC-KO) mice and *CD4-Cre Il27Ra^fl/fl^*(T-rKO) mice were described previously ^35^. Treg cell-specific deletion of IL-27p28 was achieved by breeding *Il27^fl/fl^* mice ^77^ to *Foxp3^Cre^* mice ^18^. All mice were bred and housed under specific pathogen-free conditions. Germ-free animal studies were done in collaboration with Dr. Hiutung Chu (University of California, San Diego) and those mice were housed in the dedicated germ-free facility equipped with flexible-film isolators. 8∼12 week old mice of both sexes were used and only *Foxp3^Cre^* WT littermates of the same gender served as controls in each experiment. All mice were maintained and handled in accordance with the Institutional Animal Care and Use Guidelines of University of California, San Diego and National Institutes of Health Guidelines for the Care and Use of Laboratory Animals and the Animal Research: Reporting In Vivo Experiments (ARRIVE) guidelines.

### Flow cytometry and antibodies

Cells were stained with Ghost Dye Red 780 (catalog 13-0865-T100; Tonbo Biosciences), followed by surface and intracellular antibody staining for CD4 (catalog 45-0042-82; Thermo Fisher Scientific); Ki67 (catalog 51-5698-82; Thermo Fisher Scientific); CD62L (catalog 12-0621-82; Thermo Fisher Scientific); CD8α(catalog 25-0081-82; Thermo Fisher Scientific); CD44 (catalog 48-0441-80; Thermo Fisher Scientific); IFNγ (catalog 25-7311-82; Thermo Fisher Scientific); IL-17A (catalog 48-7177-82; Thermo Fisher Scientific); CD25 (catalog 12-1522-82; eBioscience); CD83 (catalog 121508; Biolegend); TCF1 (6709S; Cell Signaling); ST2 (catalog 46-9335-82; eBioscience); and Foxp3 (catalog 53-5773-82; Thermo Fisher Scientific) at the manufacturer’s recommended concentrations. Fixation and permeabilization of cells were performed with reagents from the Tonbo Biosciences FOXP3/ Transcription Factor Staining Kit (catalog TNB-0607). To detect cytokine production, cells were stimulated in a 96-well plate with 50 ng/ml PMA, 0.5 μg/ml ionomycin, and 1 μg/ml brefeldin A (all from Sigma-Aldrich) in complete 5% RPMI media for 4 hours at 37°C before staining. An LSRFortessa or LSRFortessa X20 cell analyzer (BD Biosciences) was used for data collection, and Flowjo software (BD Biosciences) was used for data analysis.

### *In vivo* activation of Treg cells

To eliminate endogenous Treg cells, *Foxp3^DTR^* mice were injected with 50μg of DT per kg body weight intraperitoneally for two consecutive days and every other day thereafter as described previously ^27^. On the second day of DT injection, 2×10^6^ CD4^+^Foxp3Thy1.1^+^ Treg cells FACS sorted from *Foxp3^Thy^*^1^*^.1^* mice were transferred intravenously. Activated CD4^+^Foxp3Thy1.1^+^ Treg cell and CD4^+^Foxp3Thy1.1^−^ Tconv cell populations as well as controlled CD4^+^Foxp3Thy1.1^+^ Treg cells and CD4^+^Foxp3Thy1.1^−^ Tconv cells were isolated from different tissues (i.e. spleen, lung and SI LP) in DT- and PBS-treated mice 10 days after Treg cell transfer, respectively. All T cell populations were first enriched by positive selection with CD4 MojoSort™ beads (Biolegend) before FACS sorting.

### Tissue preparation and cell isolation

Spleen and lymph nodes were mechanically dissociated between frosted glass slides or with the back of a syringe plunger and filtered through a 100-μm nylon mesh to yield single-cell suspensions. For isolation of lymphocytes from lung, SI LP, LI LP or tumor, after perfusion, tissues were harvested and minced before transferring to conical tubes. The minced pieces were resuspended in 10ml of complete RPMI-1640 containing 1% P/S, 20mM HEPES, 0.05 mg/ml Liberase TL (Roche), 0.05% DNaseI (Roche) and shaken at 200 rpm for 30 min at 37°C in 50-ml Falcon tubes. The tissue suspension was collected and passed through a 70 μm cell strainer and the cells were pelleted by centrifugation at 1200 rpm. The cells were then resuspended and purified by 47% Percoll and centrifuged at 1500 rpm for 10 min. The pellet were collected, washed and resuspended in complete RPMI media.

### Gene-expression profiling

Poly-A RNA-seq was performed using 3 biological replicates for each sorted cell population. Reads were mapped to mouse genome version mm9 with STAR aligner, counts were generated using htseq/0.6.1, and differential gene expression analysis was conducted using DESeq2/1.30.1 in R. Differentially expressed genes in Treg cells compared to Tconv cells from their respective tissue origin and treatment condition were generated in DESeq2 with Negative Binomial Generalized Linear Model fitting using the Wald test for significance and Benjamini-Hochberg correction for multiple testing. The differentially expressed genes with adjusted p values > 0.05 were plotted in scatter plots and used to make the venn diagrams. Genes annotated as upregulated or downregulated in the scatterplots were used for Gene Ontology analysis, which was conducted for biological processes using enrichGO in the clusterProfiler package with the following parameters: pvalueCutoff = 0.05, qvalueCutoff = 0.02, pAdjustMethod = Benjamini-Hochberg, dropGO level 5 and simplify cutoff 0.5. Count data was transformed with variance stabilizing transformation for visualization with heatmaps and PCA plots. Violin plots were made for specific genes using normalized counts from DESeq2.

For scRNA-seq analysis, CD45^+^ immune cells isolated from the large intestine of uninfected mice and mice 10 days after *C. rodentium* infection were sent for single-cell libraries preparation according to the protocol for 10x Genomics for Single Cell 5’ Gene Expression. About 10,000 sorted CD45^+^ cells were loaded and partitioned into Gel Bead In-Emulsions. The fastq files were aligned to the mm10 mouse genome using the Cell Ranger (7.0.0) pipeline, including intronic reads. The Seurat (4.1.0) package in R (4.1.2) was used for the gene expression analysis. Cells that were dying were first removed by filtering out cells with less than 200 genes, 500 transcripts, or greater than 10% mitochondrial content. All the cells captured were then clustered and the cluster that had the highest expression of Foxp3 as well as a transcriptomic signature of Treg cells was selected for further analysis. This Treg subset included 242 cells from the infected group and 282 cells from the uninfected group. Differential gene expression analysis was done using the FindMarkers function within the Seurat package which uses the Wilcoxon Rank Sum Test. RNA-seq and scRNA-seq data are available from NCBI under accession no. GSE217949.

### qPCR analysis

For quantification of *Il27*, *Ebi3*, *Il12a*, and *Il10* expression, Tconv cells and Treg cells in different tissues from DT- and PBS-treated *Foxp3^DTR^* mice or from Treg-KO and WT littermates were sorted on a FACSAria Fusion cell sorter (BD Biosciences) with a purity of greater than 95%. For certain experiments, WT *Foxp3^YFPcre^* mice were infected with *C. rodentium* as described below. 10 days after infection, splenocytes and colonic immune cells were extracted followed FACS isolation of CD83^+^CD62L^lo^, CD83^-^CD62L^lo^ and CD83^-^CD62L^hi^ CD4^+^Foxp3YFP^+^ Treg cells. Cells were stimulated with LPS (1 μg/ml) for 6 hours at 37°C followed by RNA isolation using a RNeasy Kit (QIAGEN). Extracted RNA was converted to cDNA with an iScript cDNA Synthesis Kit (Bio-Rad), followed by qPCR reactions using SYBR Select Master Mix (Thermo Fisher Scientific). All real-time reactions were run on a 7900HT Fast Real-Time PCR System (Thermo Fisher Scientific) with the following primers: *Il27*: 5’-CTGAATCTCGATTGCCAGGAGTGA-3’ (forward) and 5’-AGCGAGGAAGCAGAGTCTCTCAGAG-3’ (reverse); *Ebi3*: 5’-CGGTGCCCTACATGCTAAAT-3’ (forward) and 5’-GCGGAGTCGGTACTTGAGAG-3’ (reverse); *Il12a*: 5’-CAGGCTACCTCCTCTTTTTG-3’ (forward) and 5’-CAGCAGTGCAGGAATAATGTT-3’ (reverse); *Il10*: 5’-CAGAGCCACATGCTCCTAGA-3’ (forward) and 5’-TGTCCAGCTGGTCCTTTGTT-3’ (reverse).

### ELISA

For quantification of the production of IL-27, IL-35 and IL-10, Tconv cells and Treg cells in different tissues from DT- and PBS-treated *Foxp3^DTR^* mice or from αCD3 mAb-treated or Citrobacter-infected Treg-KO and WT littermates were sorted on a FACSAria Fusion cell sorter (BD Biosciences) with a purity of greater than 95%. Cells were stimulated with LPS (0.5 or 1 μg/ml) for 48 or 72 hours at 37°C. Supernatant were collected and measured by ELISA kits according to the manufacturer’s instructions (catalog 438707, 440507, 431414; BioLegend). Absorbance was measured at 450 nm with a microplate reader (Molecular Devices).

### *In vitro* suppression assay

CFSE-labeled 4×10^4^ naïve CD4^+^CD25^−^CD62L^hi^ T cells from Ly5.1^+^ B6 mice and CD4^+^Foxp3YFP^+^ Treg cells in SI LP or spleen from Treg-KO mice or WT control littermates were co-cultured at the indicated ratios and stimulated with 1 μg/ml αCD3 mAb in the presence of 15×10^4^ Mitomycin C-treated T cell-depleted splenocytes for 72 hours at 37°C. CFSE dilution was assessed by FACS analysis.

### Adoptive T cell transfer study

1.6×10^6^ CD4^+^ T cells isolated from Ly5.1^+^ *Foxp3^KO^* mice mixed with 4×10^5^ CD4^+^Foxp3YFP^+^ Treg cells from Treg-KO mice or their WT littermates were intraperitoneally injected into *Rag1^−/−^* recipients. Mice were sacrificed three weeks after cell transfer or when mice reached less than 80% of their original body weight. Colonic immune cells were isolated for FACS analysis as described above.

### αCD3 mAb-induced intestinal inflammation

αCD3 mAb (Clone #: 2C11; Bio-X-Cell) were injected intraperitoneally three times (20, 20, and 20 μg per mouse) every other day. On Day 5, mice were taken down for histology, tissue preparation, cell isolation and immune staining.

### *T. gondii* Infection

The ME-49 strain of *T. gondii* was maintained in Swiss Webster and CBA/CaJ mice and tissue cysts from the brain were used for infection as previously described ^35^. For all studies, 8- to 12- week old or 6 months old mice were infected with 40 cysts of ME-49 by oral route and analyzed for parasite burden. To quantify parasite burden, qPCR was performed for DNA isolated from duodenum and liver of infected mice using primers 5’-TCCCCTCTGCTGGCGAAAAGT-3’ (forward) and 5’-AGCGTTCGTGGTCAACTATCGATTG-3’ (reverse) to determine the relative abundance of *T.gondii B1* gene to mouse *Gapdh* gene. PCR reaction was run using the standard setting on the Applied Biosystems 7900 as described previously ^35^.

### C. rodentium Infection

For infections, *C. rodentium* (DBS100 strain) were overnight cultured from a single colony in LB with nalidixic acid (Nal) from Day-1. On Day 0, each mouse was infected with 5.0 x 10^9^ CFU/mouse in a volume of 100 μl by oral gavage. On Day 10, mice were taken down for tissue preparation, cell isolation and immune staining. To quantify bacteria burden, fecal samples were also collected at day 10 post infection, the pellets were weighed in 1 mL of PBS and homogenized for 10 min at room temperature. The numbers of bacteria were counted by plating dilutions of the excess inoculums sample onto LB agar with Nal as previously described ^78^.

### Histology

To assess immunopathology, different tissues were harvested and immediately fixed in 10% formalin solution. Paraffin-embedded sections were cut (5 mm thickness) and stained with H&E as described previously ^79^. All slides were digitized and imaged using the Olympus Nanozoomer and Digital Pathology viewing software (Nikon).

### AOM/DSS-induced colitis-associated cancer

For induction of colon cancer, mice were intraperitoneally injected with 10 mg/kg AOM (Sigma-Aldrich). After 5 days, mice were supplied with 2% DSS solution for 5 days followed with normal drinking water for 15 days. The DSS cycle was repeated three times, and mice were taken down after the last DSS cycle for tissue preparation, cell isolation and immune staining ^80^.

### Statistics

Unpaired, 2-tailed Student’s t test or one-way ANOVA for multiple sample (>2) comparison were performed using GraphPad Prism 8 software (GraphPad Software).

## Author contributions

C.-H.L. and L.-F.L. conceived and designed the project. C.-H.L., C.-J.W., S.C., R.R.G., and C.-Y.H. performed the experiments. C.-H.L., R.P., E.I., J.B., M.N., M.C., R.A.M., S.A.P., H.G.D., and L.-F.L. analyzed the data. L.-L.L., M.-C.C., H.C., M.R., and J.T.C. contributed critical reagents, materials and analytic tools. C.-H.L. and L.-F.L. wrote the manuscript.

## Acknowledgments

This work was supported by NIH grants AI108651, AI127751, AI163813 (to L.-F.L.), DK110534, DK120515 (to H.C.), AI126277, AI145325, AI154644 (to M.R.), and AI132122, BX005106 (to J.T.C.). Work in M.R. lab is also supported by the Chiba University-UCSD Center for Mucosal Immunology, Allergy, and Vaccines. M.R. holds an Investigator in the Pathogenesis of Infectious Disease Award from the Burroughs Wellcome Fund. R.P. is a BioLegend fellow. R.R.G. is partly supported by a fellowship from the Crohn’s and Colitis Foundation. We thank all members of our laboratory for discussions. L.-F.L. is a scientific advisor for Elixiron Immunotherapeutics and receiving research grants from AstraZeneca and Avidity Biosciences.

