## Supplemental Figures for "Selective IL-27 production by intestinal regulatory T cells permits gut-specific regulation of Th17 immunity"

### Slide 1
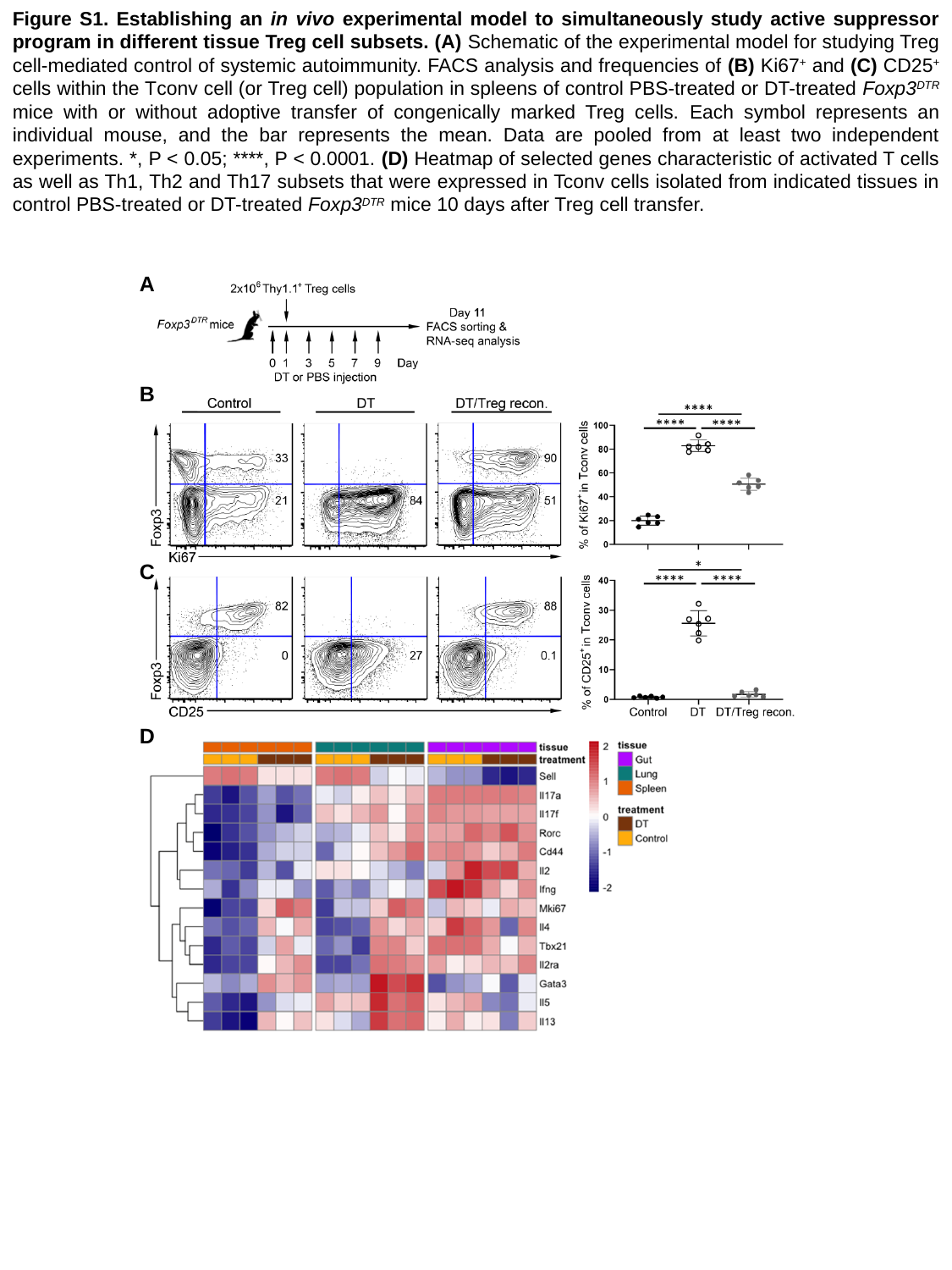

Figure S1. Establishing an in vivo experimental model to simultaneously study active suppressor program in different tissue Treg cell subsets. (A) Schematic of the experimental model for studying Treg cell-mediated control of systemic autoimmunity. FACS analysis and frequencies of (B) Ki67^+^ and (C) CD25^+^ cells within the Tconv cell (or Treg cell) population in spleens of control PBS-treated or DT-treated Foxp3^DTR^ mice with or without adoptive transfer of congenically marked Treg cells. Each symbol represents an individual mouse, and the bar represents the mean. Data are pooled from at least two independent experiments. *, P < 0.05; ****, P < 0.0001. (D) Heatmap of selected genes characteristic of activated T cells as well as Th1, Th2 and Th17 subsets that were expressed in Tconv cells isolated from indicated tissues in control PBS-treated or DT-treated Foxp3^DTR^ mice 10 days after Treg cell transfer.
A
B
C
D

### Slide 2
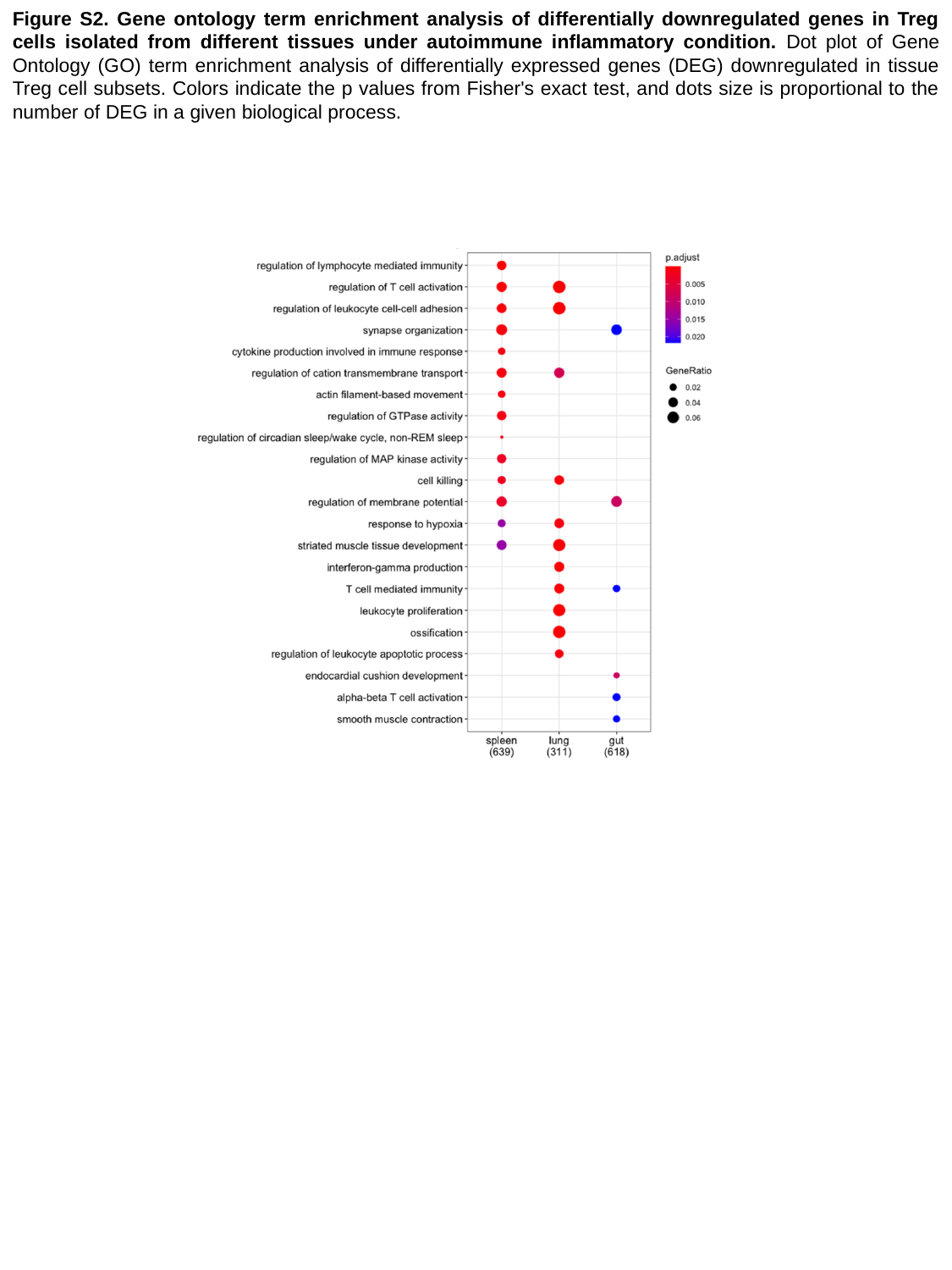

Figure S2. Gene ontology term enrichment analysis of differentially downregulated genes in Treg cells isolated from different tissues under autoimmune inflammatory condition. Dot plot of Gene Ontology (GO) term enrichment analysis of differentially expressed genes (DEG) downregulated in tissue Treg cell subsets. Colors indicate the p values from Fisher's exact test, and dots size is proportional to the number of DEG in a given biological process.

### Slide 3
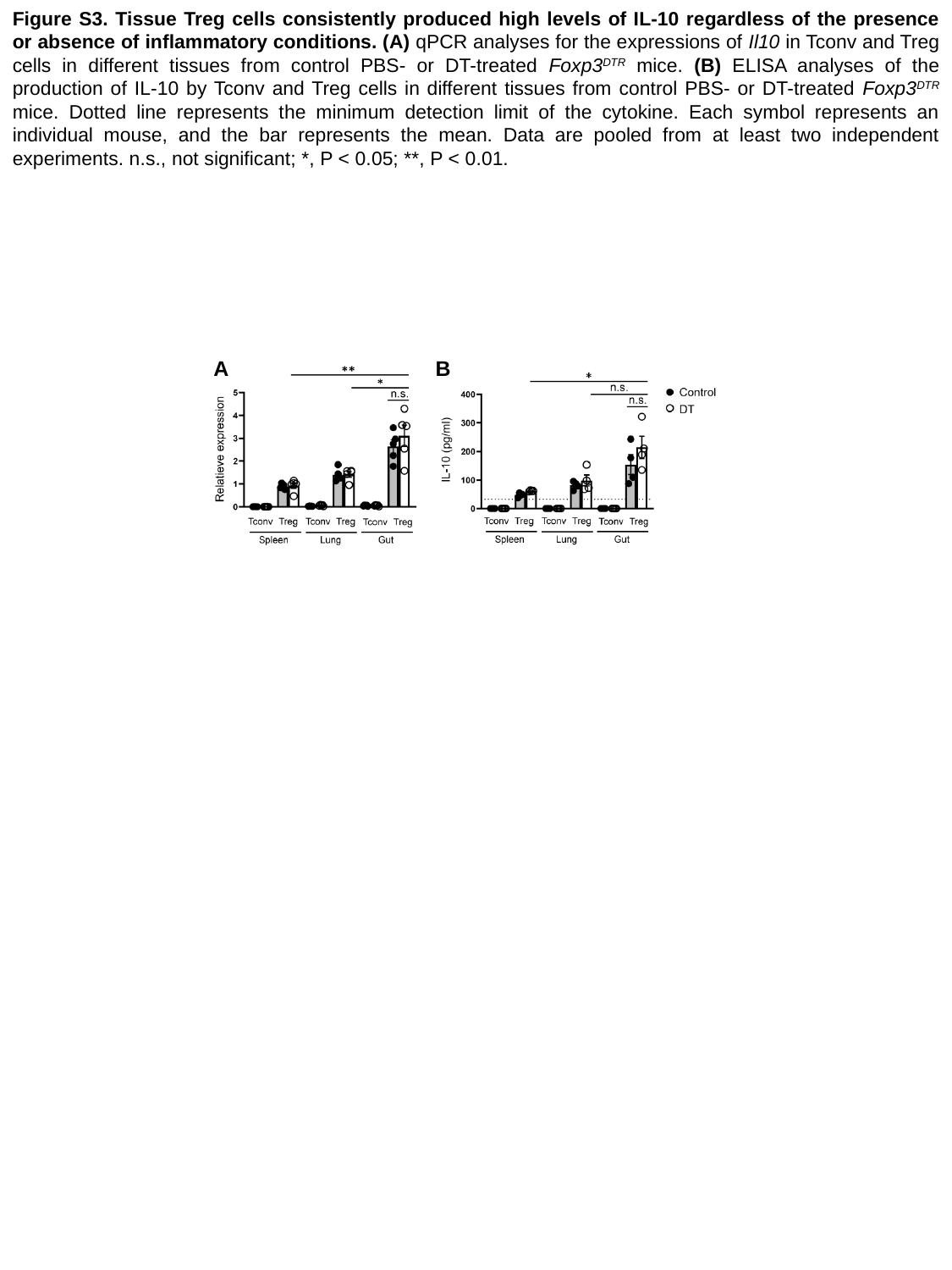

Figure S3. Tissue Treg cells consistently produced high levels of IL-10 regardless of the presence or absence of inflammatory conditions. (A) qPCR analyses for the expressions of Il10 in Tconv and Treg cells in different tissues from control PBS- or DT-treated Foxp3^DTR^ mice. (B) ELISA analyses of the production of IL-10 by Tconv and Treg cells in different tissues from control PBS- or DT-treated Foxp3^DTR^ mice. Dotted line represents the minimum detection limit of the cytokine. Each symbol represents an individual mouse, and the bar represents the mean. Data are pooled from at least two independent experiments. n.s., not significant; *, P < 0.05; **, P < 0.01.
A B

### Slide 4
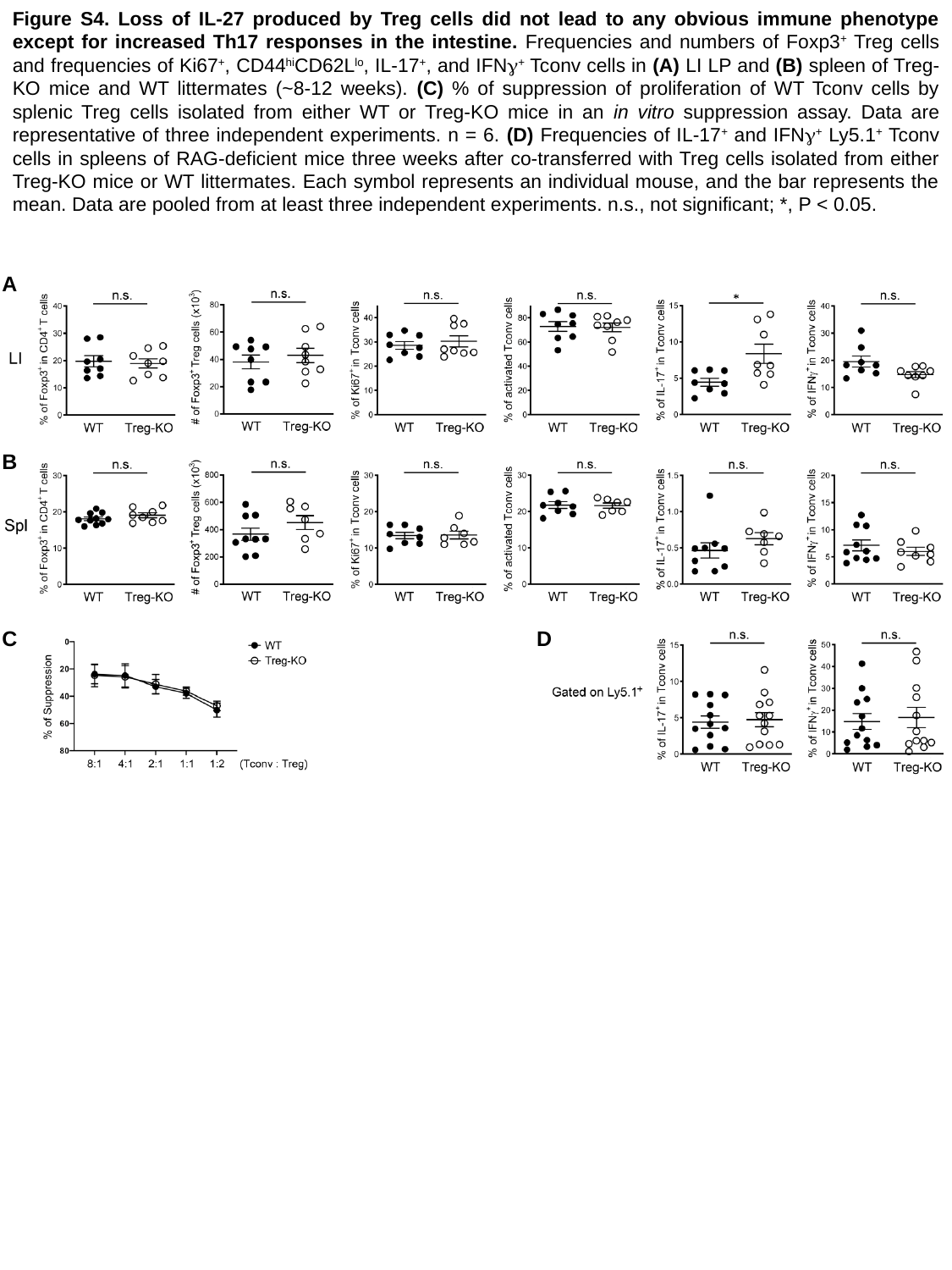

Figure S4. Loss of IL-27 produced by Treg cells did not lead to any obvious immune phenotype except for increased Th17 responses in the intestine. Frequencies and numbers of Foxp3^+^ Treg cells and frequencies of Ki67^+^, CD44^hi^CD62L^lo^, IL-17^+^, and IFNγ^+^ Tconv cells in (A) LI LP and (B) spleen of Treg-KO mice and WT littermates (~8-12 weeks). (C) % of suppression of proliferation of WT Tconv cells by splenic Treg cells isolated from either WT or Treg-KO mice in an in vitro suppression assay. Data are representative of three independent experiments. n = 6. (D) Frequencies of IL-17^+^ and IFNγ^+^ Ly5.1^+^ Tconv cells in spleens of RAG-deficient mice three weeks after co-transferred with Treg cells isolated from either Treg-KO mice or WT littermates. Each symbol represents an individual mouse, and the bar represents the mean. Data are pooled from at least three independent experiments. n.s., not significant; *, P < 0.05.
A
B
C D

### Slide 5
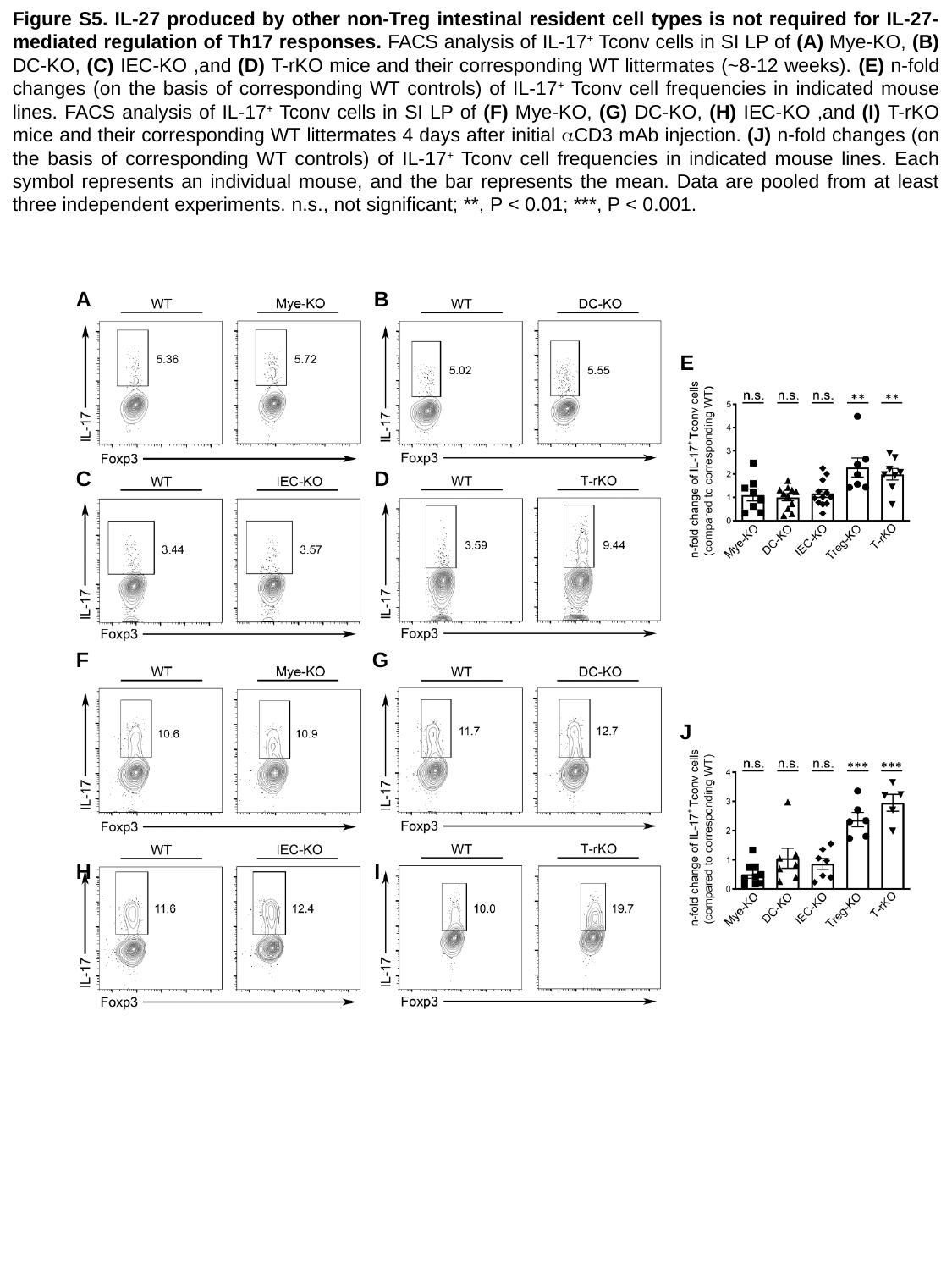

Figure S5. IL-27 produced by other non-Treg intestinal resident cell types is not required for IL-27-mediated regulation of Th17 responses. FACS analysis of IL-17^+^ Tconv cells in SI LP of (A) Mye-KO, (B) DC-KO, (C) IEC-KO ,and (D) T-rKO mice and their corresponding WT littermates (~8-12 weeks). (E) n-fold changes (on the basis of corresponding WT controls) of IL-17^+^ Tconv cell frequencies in indicated mouse lines. FACS analysis of IL-17^+^ Tconv cells in SI LP of (F) Mye-KO, (G) DC-KO, (H) IEC-KO ,and (I) T-rKO mice and their corresponding WT littermates 4 days after initial αCD3 mAb injection. (J) n-fold changes (on the basis of corresponding WT controls) of IL-17^+^ Tconv cell frequencies in indicated mouse lines. Each symbol represents an individual mouse, and the bar represents the mean. Data are pooled from at least three independent experiments. n.s., not significant; **, P < 0.01; ***, P < 0.001.
A B
 E
C D
F G
 J
H I

### Slide 6
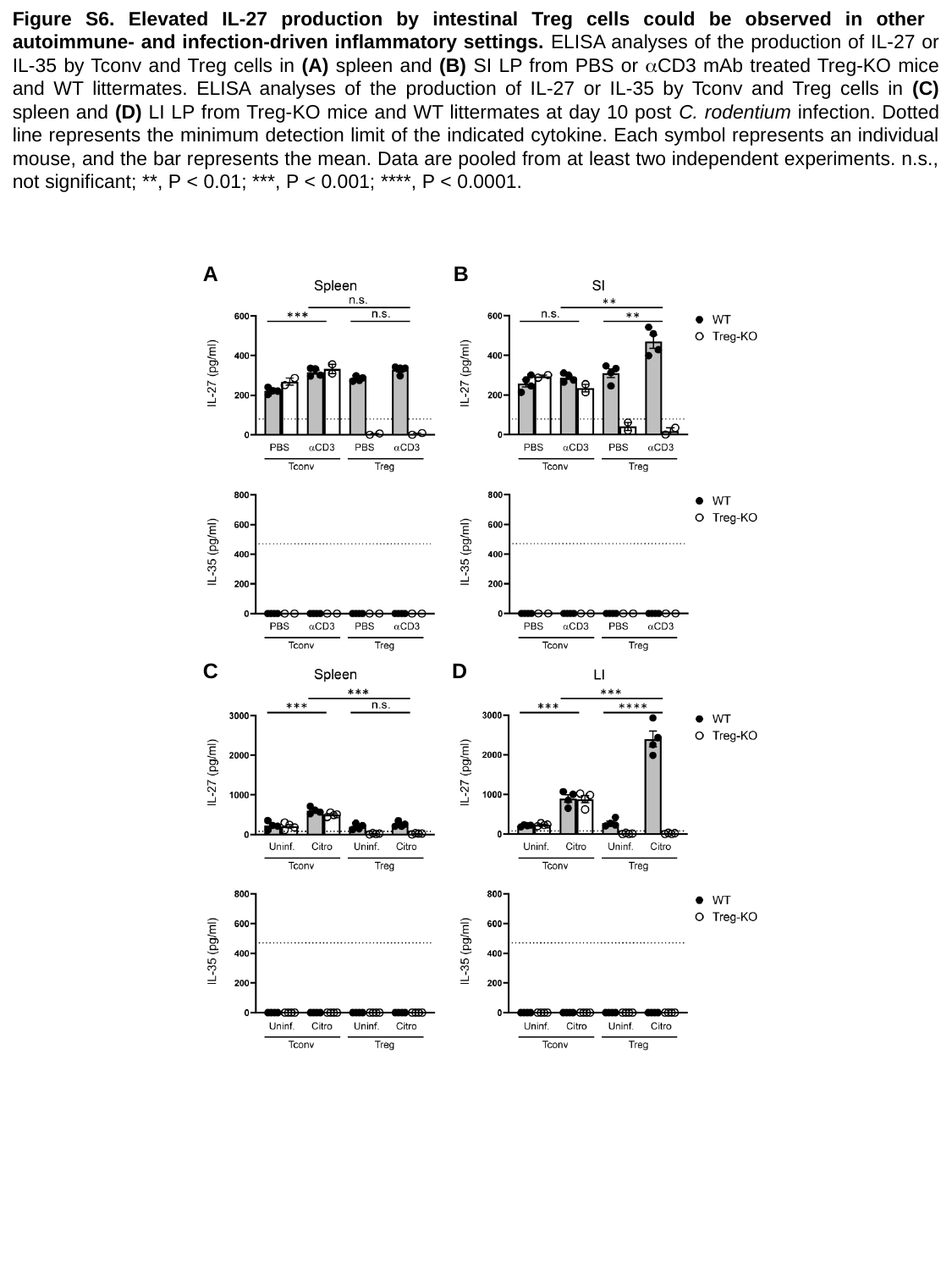

Figure S6. Elevated IL-27 production by intestinal Treg cells could be observed in other autoimmune- and infection-driven inflammatory settings. ELISA analyses of the production of IL-27 or IL-35 by Tconv and Treg cells in (A) spleen and (B) SI LP from PBS or αCD3 mAb treated Treg-KO mice and WT littermates. ELISA analyses of the production of IL-27 or IL-35 by Tconv and Treg cells in (C) spleen and (D) LI LP from Treg-KO mice and WT littermates at day 10 post C. rodentium infection. Dotted line represents the minimum detection limit of the indicated cytokine. Each symbol represents an individual mouse, and the bar represents the mean. Data are pooled from at least two independent experiments. n.s., not significant; **, P < 0.01; ***, P < 0.001; ****, P < 0.0001.
A B
C D

### Slide 7
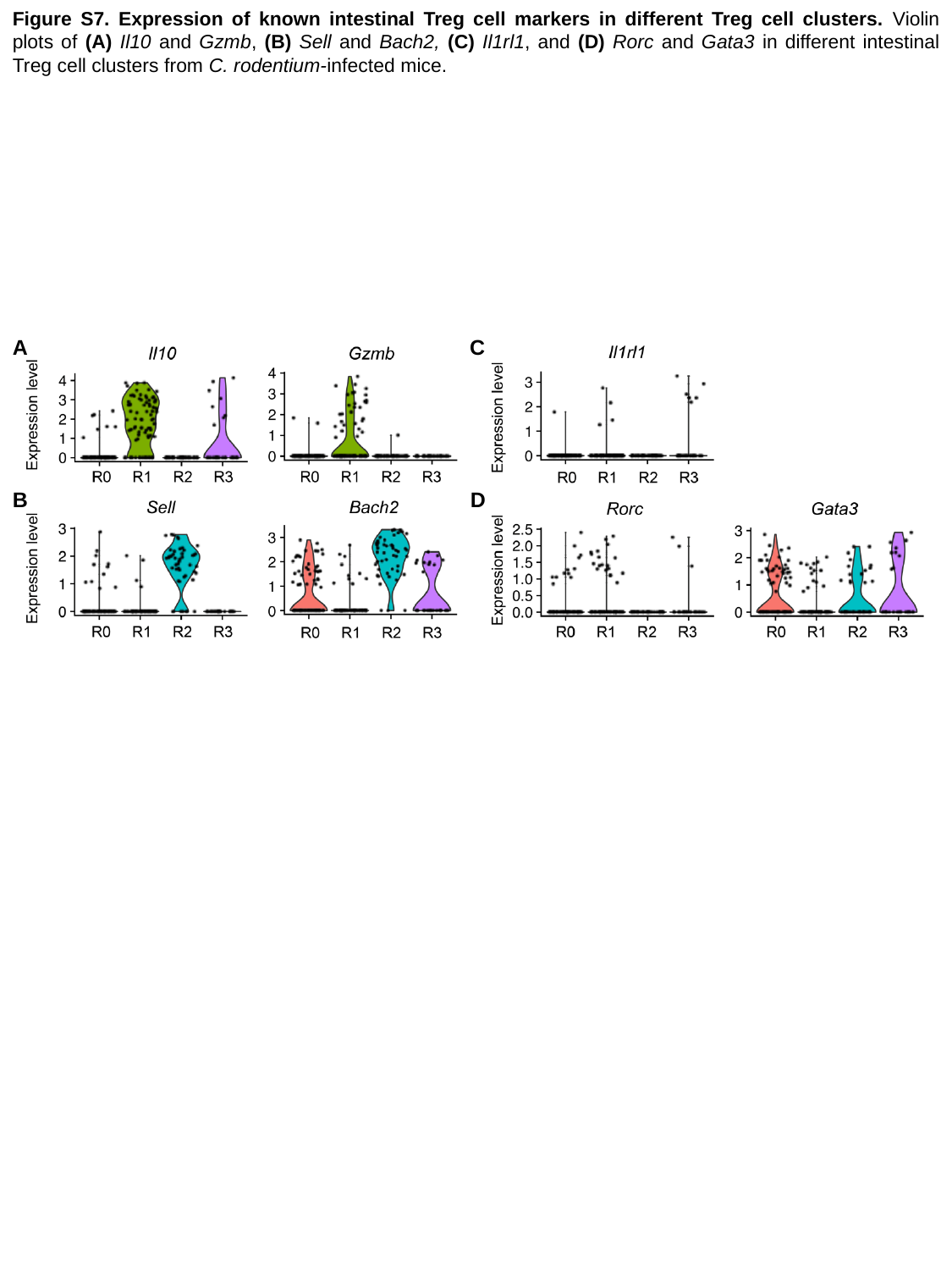

Figure S7. Expression of known intestinal Treg cell markers in different Treg cell clusters. Violin plots of (A) Il10 and Gzmb, (B) Sell and Bach2, (C) Il1rl1, and (D) Rorc and Gata3 in different intestinal Treg cell clusters from C. rodentium-infected mice.
A C
B D
